# Accurate and robust inference of genetic ancestry from cancer-derived molecular data across genomic platforms

**DOI:** 10.1101/2022.02.01.478737

**Authors:** Pascal Belleau, Astrid Deschênes, David A. Tuveson, Alexander Krasnitz

## Abstract

Genetic ancestry-oriented cancer research requires the ability to perform accurate and robust genetic ancestry inference from existing cancer-derived data, including whole exomes, transcriptomes and targeted gene panels, very often in the absence of matching cancer-free genomic data. Here we examine the feasibility and accuracy of such computation. In order to optimize and assess the performance of the ancestry inference for any given input cancer-derived molecular profile, we have developed a data synthesis framework. In its core procedure, the ancestral background of the profiled patient is replaced with one of any number of individuals with known ancestry. Data synthesis is applicable to multiple profiling platforms and makes it possible to assess the performance of inference specifically for a given molecular profile, and separately for each continental-level ancestry. This ability extends to all ancestries, including those without statistically sufficient representation in the existing cancer data. We further show that our inference procedure is accurate and robust in a wide range of sequencing depths. Testing our approach for three representative cancer types, and across three molecular profiling modalities, we demonstrate that global, continental-level ancestry of the patient can be inferred with high accuracy, as quantified by its agreement with the golden standard of the ancestry derived from matching cancer-free molecular data. Our study demonstrates that vast amounts of existing cancer-derived molecular data potentially are amenable to ancestry-oriented studies of the disease, without recourse to matching cancer-free genomes or patients’ self-identification by ancestry.

## Introduction

There is ample epidemiological evidence that race and/or ethnicity are important determinants of incidence, clinical course and outcome in multiple types of cancer [1–5]. As such, these categories must be taken into account in the analysis of molecular data derived from cancer. A number of recently published large-scale genomic studies of cancer point to differences in the molecular make-up of the disease among groups of different ancestral background and to the need for more molecular data to power discovery of such differences [6–11].

Ancestry annotation of cancer-derived data largely draws on two sources. One is a patient’s self-identified race and/or ethnicity (SIRE). SIRE is often missing, sometimes inaccurate and usually incomplete. As a recent analysis [12] of PubMed database entries since 2010 reveals, patients’ SIRE is massively under-reported in genome and exome sequencing studies of cancer, with only 37% of these reporting race, and 17% reporting ethnicity. Furthermore, SIRE is not always consistent with genetic ancestry. Finally, a self-declaring patient is often given a choice from a small number of broad racial or ethnic categories, which fail to capture complete ancestral information, especially in cases of mixed ancestry [13].

A far more accurate and detailed ancestral characterization may be obtained by genotyping a patient’s DNA from a cancer-free tissue. Powerful methods exist for ancestry inference from germline DNA sequence [14–17]. These methods were recently used to determine ancestry of approximately 10,000 patients profiled by The Cancer Genome Atlas (TCGA) [7, 8]. However, genotyping of DNA from patient-matched cancer-free specimens is not part of standard clinical practice, where the purpose of DNA profiling is often identification of mutations with known oncogenic effects, such as those in the Catalog Of Somatic Mutations In Cancer (COSMIC) database [18]. As a result, it is not performed routinely outside academic clinical centers or major research projects. There also are studies yielding sequence data from tumors, whose purpose does not require germline profiling. RNA sequencing (RNA-seq) for expression quantification is in this category. Finally, peripheral blood is most often the source of germline DNA in the clinic, but this is not always the case for diseases of the hematopoietic system, such as leukemia, wherein cancer cells are massively present in circulation. In summary, matched germline DNA sequence is not universally available for cancer-derived molecular data. In such cases, it is necessary to infer ancestry from the nucleic acid sequence of the tumor itself.

Standard methods of ancestry inference commonly rely on population specificity of germline single-nucleotide variants (SNV). Whole-genome (WGS) or whole-exome sequences (WES), at depths sufficient for reliably calling single-nucleotide variants, and readouts from genotyping microarrays, are therefore data types most suitable for this purpose. However, such detailed DNA profiling is often not performed in molecular studies of cancer. In such cases, it is necessary to infer ancestry from other types of tumor-derived data, including RNA sequence and DNA sequence for a small panel of genes, e.g., FoundationOne^®^ CDx [19].

For all types of tumor-derived sequence, accurate inference of ancestry is a potential challenge. Tumor genome is often replete with somatic alterations, including loss of heterozygosity (LOH), copy number variants (CNV), translocations, microsatellite instabilities and SNV. Of these, structural variants, especially LOH and CNV, are the most likely to affect the genetic ancestry calls, but other types of of alterations also are, to various degrees, potential obstacles to accurate ancestry inference. Tumor RNA-seq presents additional challenges, namely, extremely uneven coverage of the transcript due to a broad range of RNA expression levels and distortions due to allele-specific expression. Gene panels represent a very small fraction of the genome, whose sufficiency for ancestry inference is not clear and may vary from panel to panel. In addition, cancer gene panels are enriched in cancer driver genes, which tend to undergo somatic alteration more frequently than other parts of the genome.

Important recent publications on ancestral effects in cancer reported patient ancestry inferred from matching cancer-free DNA [7, 8, 11]. At the same time, there has been much less work on ancestry inference from tumor-derived nucleic acids. A recent analysis of tumor genomes from TCGA and GEO repositories, profiled by SNP microarrays, demonstrated a high degree of coincidence between patient ancestries inferred from these data and those inferred from SNP profiles of matching germline genomes [20]. This study did not report inference results from other molecular profiling modalities. Similar agreement has been found, for a set of over 300 cancer cell lines, between the self-declared race/ethnicity of the donors and ancestry inferred from the SNP array data [8], but that finding was not validated against matching cancer-free data. Ancestry was also inferred in two large collections of cancer cell lines using SNP microarray data [21, 22]. In the absence of matching cancer-free genotypes or self-declared ancestry of the donor the inference accuracy could not be assessed in these two studies. Ancestry inference from RNA sequences, 174 of which were derived from cancer tissue specimens, was considered in a recent study [23]. However, these inferred ancestries were neither compared to ancestry calls from germline sequence nor to self-declared ancestries for accuracy assessment. Ancestry has been inferred for a large set of patient cases profiled with the FoundationOne^®^ CDx gene panel [19], but these ancestry calls were neither compared to those from the germline sequence nor to the patients’ SIRE. A more recent study compared, with encouraging results, ancestry inference from cancer-derived FoundationOne^®^ CDx data to matching cancer-free ancestry calls, but this analysis was confined to lung cancer in mixed American super-population. To our knowledge, no systematic computational framework for ancestry inference from cancer-derived molecular data, across assay and cancer types, has been developed to date. There is presently no ability to assess the inference accuracy specifically for a given input tumor-derived molecular profile with all its attendant properties, including the data quality and the depth of coverage. Reliable and accurate ancestry inference from tumor-derived nucleic acids thus represents an unmet need, which the present work aims to address.

For this purpose, we designed an inference procedure having in mind a scenario, likely to occur in studies of existing data or of archived tissue specimens, with an input molecular profile of a tumor from a single patient, and no matching cancer-free sequence available. The profile in question may have its unique set of sequence properties. These include the target sequence and uniformity of its coverage, depth, read length and sequencing quality. These profile-specific properties may be vastly dissimilar from those in the available public data sets with reliably known genetic ancestry of the patients. Furthermore, not all ancestries are equally easy to infer: for example, a Mixed American ancestral category is sometimes difficult to distinguish either from African or from European ancestry. This profile specificity would make it impossible to confidently assess the accuracy of the inference procedure for the input profile from its performance with the public cancer-derived data in aggregate. In order to overcome this difficulty, we develop a computational technique, which is described schematically in ***Figure 1*** wherein the ancestral background of the patient is supplanted in the input profile by one of an unrelated individual with known ancestry. We next apply established methods of ancestry inference to this synthetic profile and compare the result to that known ancestry. Generating multiple such synthetic profiles allows us to assess how accurate the ancestry inference is for the patient, both overall and as a function of the profile’s continental-level ancestry. Furthermore, using synthetic data, we are able optimize the inference procedure with respect to parameters on which it depends. Importantly, this assessment and optimization procedure does not require the profile in question to be part of a larger data set from a cohort of patients with a similar diagnosis. Very often in public cancer-derived data, such cohorts do not provide statistically meaningful representation of non-European ancestries. This insufficiency is not an impediment to the application of our methodology.

**Figure 1.**
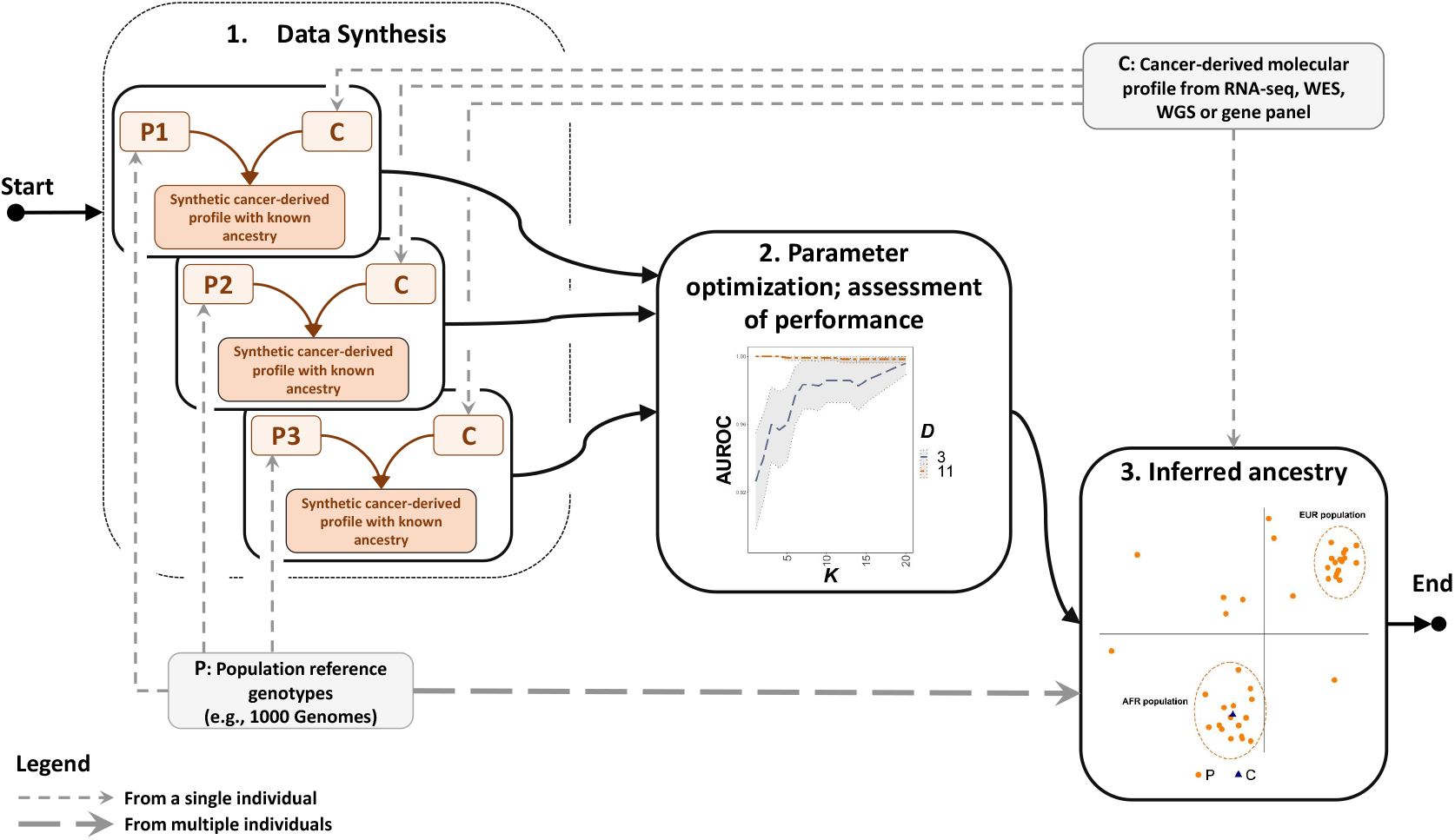
An overview of genetic ancestry inference from cancer-derived molecular data using data synthesis.

In the following, we assess the accuracy of global ancestry calls from tumor exomes, narrowly targeted gene panels and RNA sequences, in comparison to such calls from matching germline genotypes, as profiled by exome sequencing or SNP microarrays. We do so for three cancer types, namely, pancreatic adenocarcinoma (PDAC) and ovarian cystadenocarcinoma (OV) as representative types of epithelial tumors, and acute myeloid leukemia (AML), as an example of hematopoietic malignancy. Each of these data sets represents a unique challenge for patients’ ancestry inference.

OV is characterized by massive copy number alterations, often spanning much of the genome. Our PDAC data originate from patient-derived organoid (PDO) models of the disease [24]. In PDO, near-100% tumor purity is achieved, exacerbating effects of copy number loss and loss of heterozygosity on the sequence. In AML the peripheral blood, the usual source of cancer-free DNA, may be severely contaminated by the cancer.

## Methods and Materials

### Data sets and pre-processing

The data sets used in this work originate from three sources: TCGA collection for ovarian cystadenocarcinoma [25], Beat AML clinical trial [26], and a study of pancreatic ductal adenocarcinoma (PDAC) using patient-derived organoids [24]. For all three, the data used are summarized, in the form of Venn diagrams and included cancer DNA (whole-exome or whole-genome) sequence, cancer RNA sequence and matching DNA (whole-exome or whole-genome) sequence. In all cases, read data mapped to the hg38 version of the human genome were used. In order to study ancestry inference from targeted panels, the cancer-derived whole-exome data were reduced to reads mapping to the FoundationOne^®^ CDx cancer-related gene panel [27]. Reads in the cancer-derived data were filtered for quality using a cutoff phred score of 20. Following this filter, single-nucleotide substitutions were called at all positions with read coverage of at least 10, using Varscan version 2.4.4 [28]. This set of positions is called the high-confidence substitution (HCS) set in the following. From the 1000 Genomes (1KG) variant call data in the Variant Call Format (VCF) [29], genomic positions where substitution variants occur at a frequency of at least 0.01 in at least one of the super-populations comprising 1KG were selected as a basis for the ancestry inference. This set is referred to as the high-frequency substitution (HFS) set in the following. At the HFS positions in the cancer-derived profile with the coverage above 10, the genotype was called. This set of positions is referred to as high-confidence genotype (HCG) set in the following. In the HCG set, the total read count and the read counts for the reference and the alternative (according to HFS) alleles were determined. A genotype at an HCG position was considered undetermined if the excess of the total read count over the sum of the reference and alternative counts was inconsistent with the error of 0.001 at the *p* = 0.001 level of significance. The same rule was used to call a heterozygous genotype. The HCG genomic positions were pruned to reduce correlation between neighboring genotypes using Bioconductor SNPRelate package version 1.22.0 [30]), resulting in the pruned high-confidence genotype (PHCG) set of positions.

### Ancestry inference

***Figure 2*** lays out the workflow for ancestry inference. For a given cancer-derived profile, principal component analysis of the 1KG genotypes reduced to the PHCG was performed, and *D* top principal components retained. The patient genotype reduced to PHCG was projected onto the subspace spanned by these *D* components. Within this subspace, the patient’s ancestry was called as that of the 1KG super-population with the highest number of 1KG individuals among *K* nearest neighbors of the patient’s genotype, using Euclidean distance in the *D*-dimensional subspace. If two or more super-populations were found tied in the nearest-neighbor count, no ancestry call was made for the patient. Only two such ties were observed in this work.

**Figure 2.**
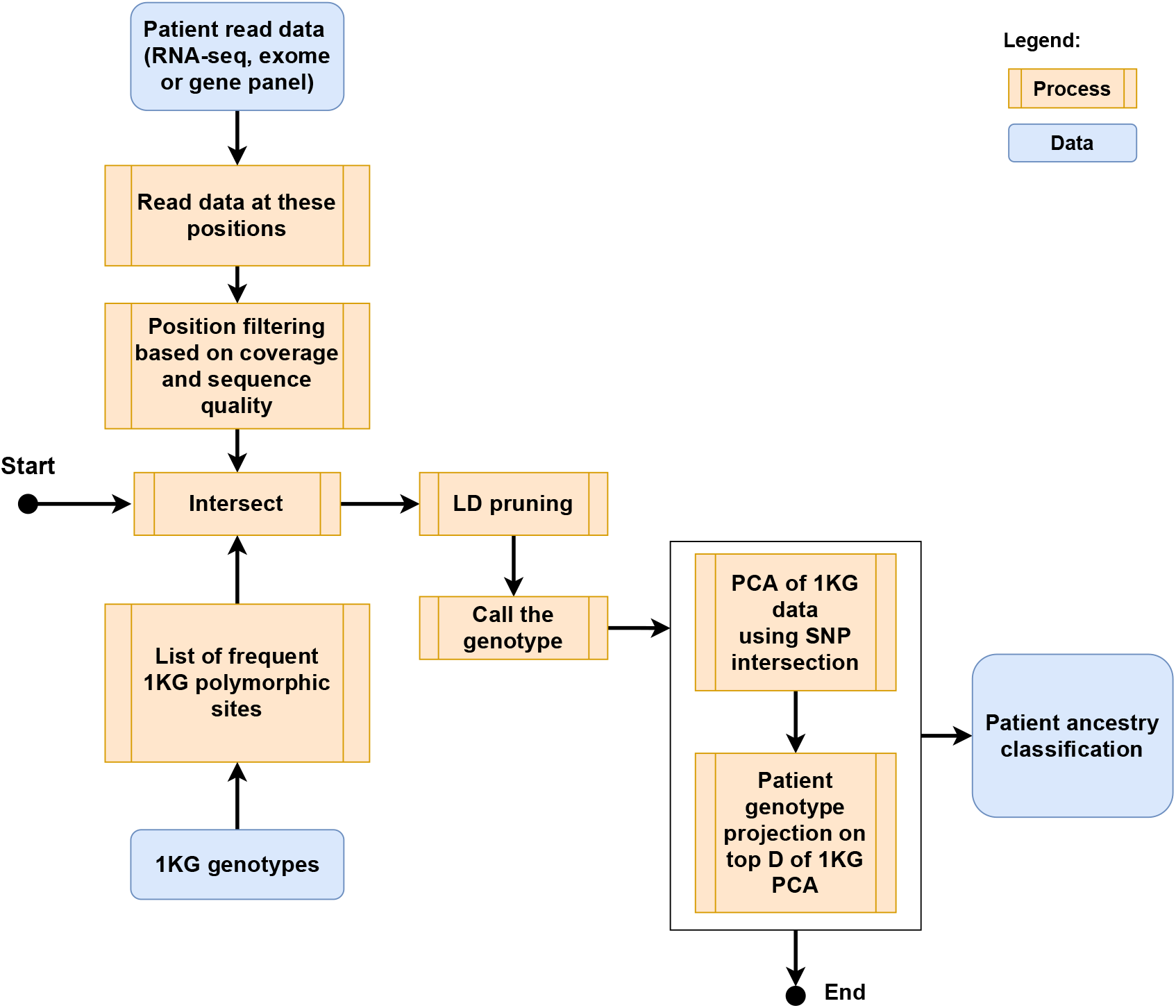
A flowchart of the inference of genetic ancestry.

### Measures of performance

We evaluate the performance of the ancestry inference by comparison to the ancestry inferred from the matching cancer-free data, wherever the latter are available. This is the case for the entirety of Beat AML and the OV data. For both, we infer the ancestry from the matching cancer-free exome profiles. In the case of OV data, we also compare the results to the consensus ancestry call [7]. In the case of PDAC matching cancer-free WGS data are available for 22 patient cases (***Figure 3***), and our assessment of accuracy is based on this subset of the data. We compute, for each dataset, the 5 × 5 confusion matrix (CM) for the 1KG superpopulation calls from the cancer-derived and cancer-free data sources. From the CM, the call accuracy is computed as the sum of the diagonal terms divided by that of the whole CM. Since the ancestral composition of all data sets considered here is heavily skewed towards the European super-population, we also compute the multi-class version of the area under the receiver operating characteristic curve (AUROC) [31]. AUROC is a measure of the call quality which compensates for the asymmetry in the class sizes.

**Figure 3.**
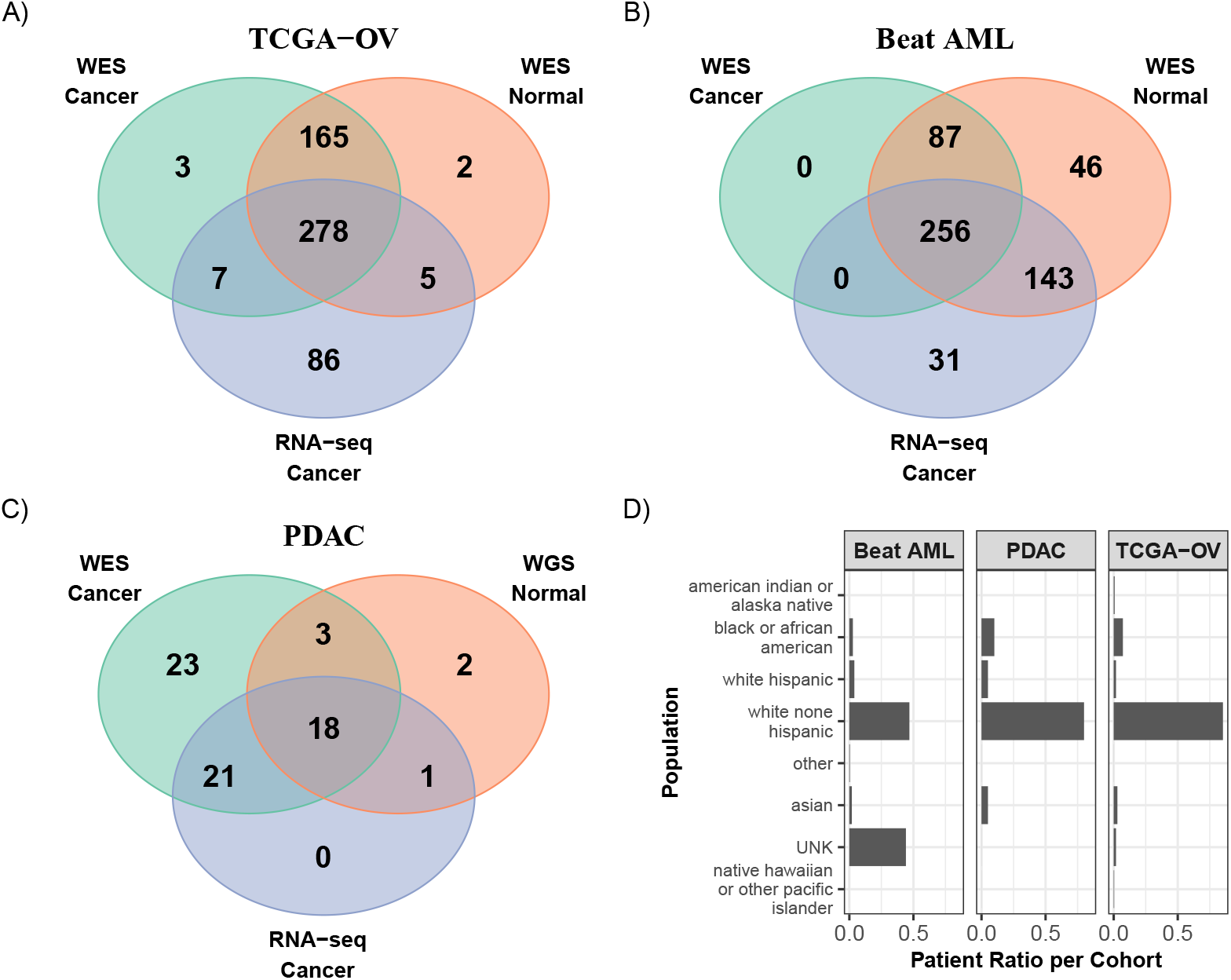
Summary of the molecular data used in this study. These originate from three patient cohorts: **A)** TCGA ovarian cancer **B)** acute myeloid leukemia and **C)** pancreatic ductal adenocarcinoma library of patient-derived organoids. **D)** The distribution of the patients by SIRE for Beat AML, PDAC and TCGA-OV cohorts. UNK means not reported or unknown.

We use an R package pROC (CRAN version 1.16.2) [32] for this purpose, and compute both the class-specific AUROC for each super-population and the 5-class AUROC. In the class-specific case, we use a version DeLong’s algorithm [33, 34] as implemented in the pROC package to compute the AUROC confidence intervals. In the 5-class case the confidence intervals are computed using bootstrap with 100-fold sampling.

### Data synthesis

Data synthesis is defined here as replacement of the sequence variants detected in a cancer-derived profile *P* by those found in the genome of an unrelated individual *U*. Ingredients required for this procedure are: (a) allele fraction (AF) estimates in *P* and (b) the haplotype of *U* in the portion of the genome covered by *P*. With this knowledge, the procedure, depicted in ***Figure 4***, consists of the following steps. First, sequence reads comprising *P* are distributed at random among the alleles with probabilities equal to the observed allele fractions. Second, in each haplotype block in the genome of *U* that is covered by *P*, allele assignment is made at random, yielding variant and reference read counts for each substitution in the genome of *U* within the scope of *P*.

**Figure 4.**
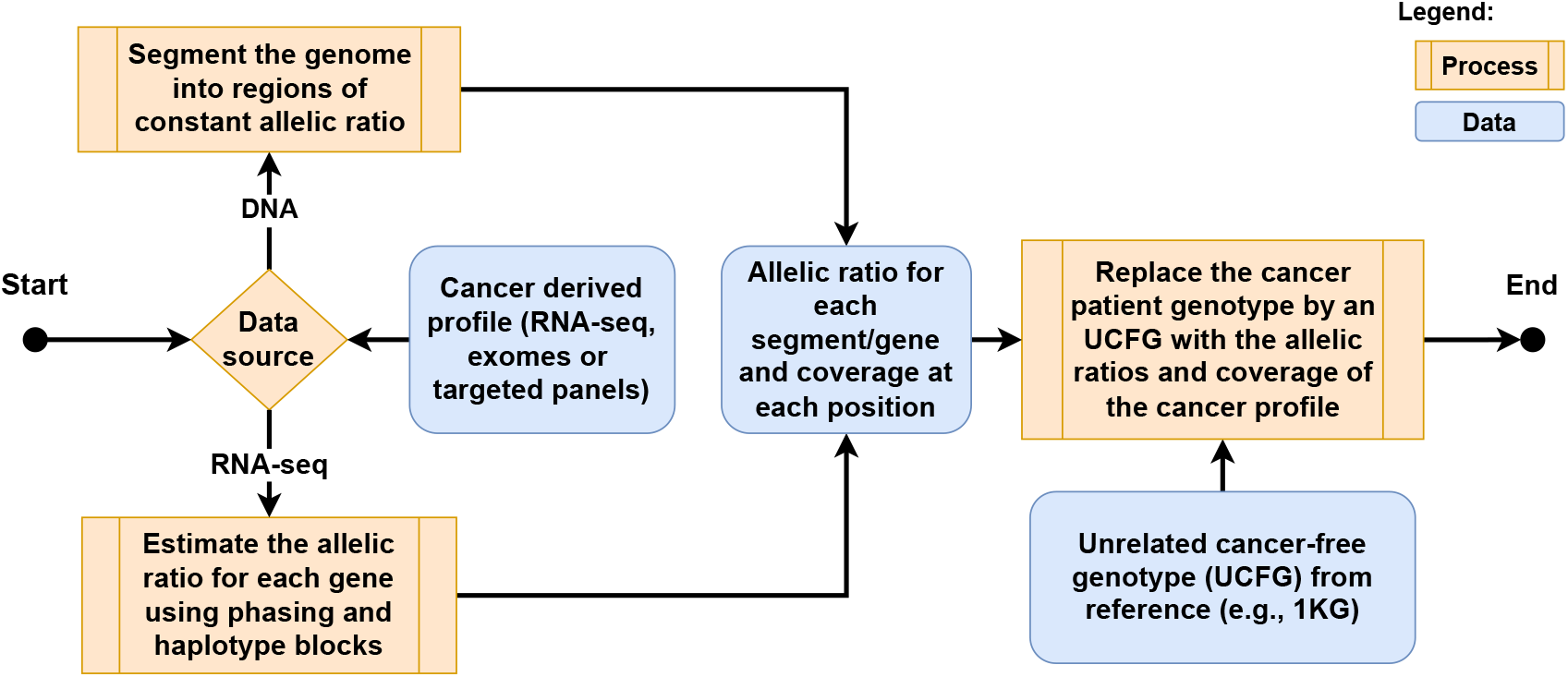
A schematic overview of the data synthesis process.

### Inference parameter optimization using synthetic data

In order to optimize ancestry inference parameters *D* and *K* for a given cancer-derived molecular profile, we generate a synthetic data set by repeatedly pairing the profile with 1KG genomes. A subset of 780 1KG genomes is set aside for this purpose by drawing at random 30 genomes from each of the 26 ancestral populations represented in 1KG Table S1. Genetic ancestry is then inferred for each of the 780 synthetic profiles following the procedure described in the Ancestry Inference sub-section, each time with the 1KG genome used for synthesis removed from the reference data set. The inference performance is then assessed as the 5-class AUROC, as explained in the Measures of Performance subsection. AUROC is computed for the *D, K* pairs in a range of values of these parameters, and the optimal *D, K* pairs yielding the highest accuracy are identified. Throughout this work, AUROC was computed for all *D* and *K* in the rectangle 3 ≤ *D* ≤ 11; 3 ≤ *K* ≤ 15. For all combinations of data sources and profiling modalities considered, a set of *D, K* pairs was found where the performance was optimal or differed from the optimum by no more than 3% (***Figure 5***).

**Figure 5.**
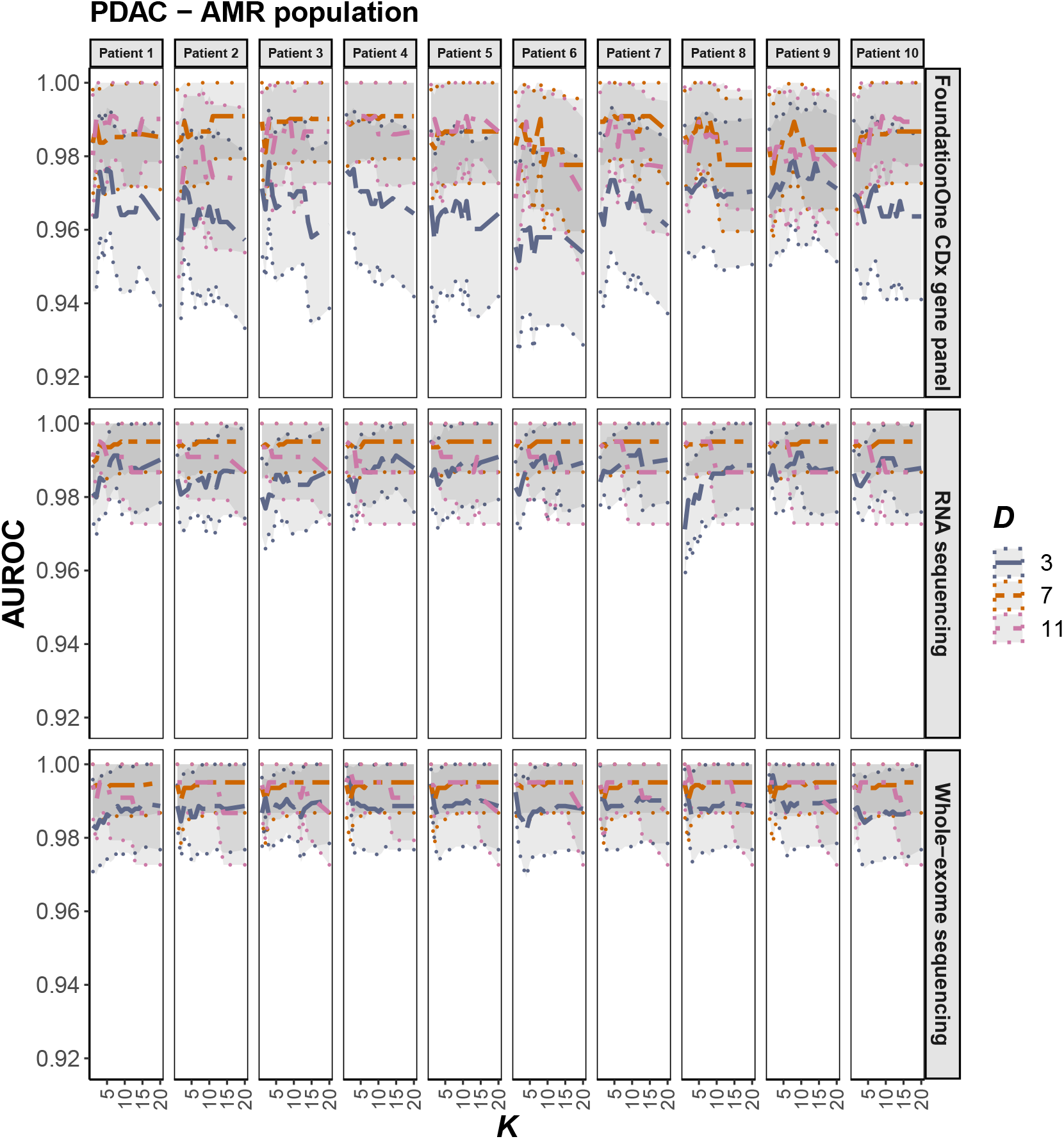
Dependence of AMR-specific AUROC on the inference parameters *D* and *K*, computed using data synthesis for 10 PDAC patients and the three profiling modalities: WES, RNA-seq and FoundationOne^®^ CDx panels. The central AUROC values are shown in solid, and the 95% CI in dashed, lines.

#### Determination of allele fractions

As the Data Synthesis subsection makes clear, knowledge of allele fractions (AF) in a cancer-derived profile is a prerequisite for data synthesis. We describe a 3-step AF estimate procedure which relies exclusively on the cancer-derived molecular profile, in the absence of a matching cancer-free genotype from the patient, as would be the case for the intended application of our methods. First (step 1), the loss-of-heterozygosity (LOH) regions are delineated. Next (step 2), the regions of allele imbalance where AF differs significantly from 1/2 are identified. Finally (step 3), AF are computed throughout the regions of allele imbalance. These steps are implemented differently, depending on whether the profile originates in the cancer DNA or RNA. We now discuss these steps, in turn for the DNA- and the RNA-derived profiles (Figure S3).

For the DNA-derived profiles, the LOH regions (step 1) are detected as follows. An LOH region in *P* must fit into a gap *G* between any two consecutive HCS positions, where all the observed genotypes are consistent with homozygosity. Any region within *G* is then considered an LOH region (see Figure S3 b) if it contains *k*_1_ PHCG positions with *k*_1_ ≥ *k*_*min*_ and for which the 1KG frequencies *F*_*i*_, 1 ≤ *i* ≤ *k*_1_ of the alleles observed in the cancer-derived profile *P* satisfy

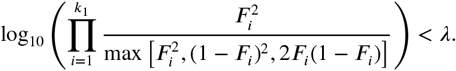

PHCG positions only are used for this purpose, to reduce correlations due to linkage. The values of *k*_*min*_ and were chosen so as to maximize, in TCGA OV data set, the overlap between the regions found to be LOH by these criteria and the published LOH regions ASCAT2 files from NCI’s Genomic Data Commons [35, 36]. The latter were determined with full knowledge of the patient’s cancer-free genotype. The optimal values were found to be *k*_*min*_ = 3 and *λ* = −3.

Step 2 is based on the notion of an “empty box” (see Figure S3 b). By this, we mean a contiguous region where the allele fraction of 1/2 is inconsistent with the read counts for the reference and alternative alleles at the HCS positions it contains. An empty box is constructed as follows. First, we consider sliding windows, each encompassing *k*_2_ consecutive HCS positions not separated by an LOH region. A window is called asymmetric if (a) for no less than *k*_2_ −1 of the positions the minor allele count is outside the inner-quartile range (IQR) of the binomial distribution with the minor AF of *f*_0_ = 1/2 and (b) satisfy

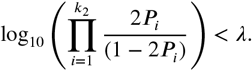

where *P*_*i*_ = P(*X*_*i*_ ≤ number of reads covering the minor allele at position *i*) and, *X*_*i*_ is the binomial distribution with the number of trials equals the coverage at the position *i* and the probability of success *p* = 1/2. In this work, *λ* = −3. A polymorphic position is called asymmetric if it belongs to at least one asymmetric window. An empty box is a region with no less than *k*_2_ polymorphic positions, all of which are asymmetric. We used *k*_2_ = 10 throughout this work.

At step 3, in the case of DNA, we consider contiguous genome regions of allele asymmetry identified at step 2. Each of these may consist of sub-regions with differing allele fractions. To detect these sub-regions, we “seed” the first sub-region with *k*_3_ HCS positions at the region’s boundary and, in this window, estimate the minor allele fraction. We consider the adjacent window *W* of *k*_3_ HCS positions *k*_3_ + 1 through 2*k*_3_ and apply to it the empty box criteria as described for step 2, with *f*_0_ set to the estimated minor allele fraction of the first window. If the criteria are satisfied, *W* becomes the seed of the next sub-region, and the process is repeated. Otherwise, HCS position *k*_3_ + 1 is added the first sub-region and *W* is shifted to start at *k*_3_ + 2, etc.

In the case of a cancer-derived RNA profile, the expressed allele fractions are, in general, gene specific. Therefore the steps 1 and 2 (condition b), as described above, are performed separately for each gene, assuming the minor allele fraction to be constant throughout the gene. Step 3 is then reduced to an empirical estimate of the minor allele fraction using read counts from all HCS positions within the gene.

### Down-sampling of sequence data

In order to down-sample the sequence data to a desired fraction *f* of the original coverage, we sampled reads from the original patient profile *P* with the Bernoulli probability *f* without replacement. The ancestry inference procedure was then performed with the resulting sample of reads.

### Software availability

An R language implementation of ancestry inference methods introduced in this work is available upon request to.

### Schematic overviews and figures

All schematic overviews were produced with draw.io version 15.7.3 (http://www.diagrams.net).

The Venn diagrams in ***Figure 3*** were produced with CRAN packages VennDiagram version 1.6.20 [37] and multipanelfigure version 2.1.2 [38].

The bar plot graph in ***Figure 3*** was produced with CRAN package ggplot2 version 3.3.5 [39].

The AUROC graphs in ***Figure 5*** was produced with CRAN packages ggplot2 version 3.3.5 [39] and cowplot version 1.1.1 [40].

## Results

We assessed the performance of genetic ancestry inference from three genomic data types: whole exomes, gene panels targeting exomes of several hundred cancer-related genes each and RNA sequences. Throughout the study, we used the 1000 Genomes (1KG) data set, with no relatives for the individuals included [41, 42], as reference, against which patient molecular data were compared to infer continental-level global ancestry. The latter is defined as a categorical variable taking five values: African (AFR), East Asian (EAS), European (EUR), Mixed American (AMR) and South Asian (SAS). These are called super-populations in the 1KG terminology. Each super-population comprises a number of subcontinental-level populations [42].

Our assessment relied on molecular data collected from three patient cohorts, each representing a cancer type, namely, tissue donors to the Cold Spring Harbor Laboratory (CSHL) pancreatic ductal adenocarcinoma (PDAC) library of patient-derived organoids; acute myeloid leukemia (AML) patients enrolled in Beat AML clinical trial; and patients comprising TCGA ovarian cancer cohort (TCGA-OV) [25]. In these cohorts, patient molecular data were available from tissue specimens both of cancer and cancer-free. ***Figure 3*** and Supplementary Table S2 contain a summary of molecular data underlying the study.

We employed principal-component analysis (PCA) as our inference tool of choice, and applied it as follows (***Figure 2***) [16].

As a basis for the analysis, we used genotypes at genomic positions where single-nucleotide sequence variants occurred with a frequency above a threshold in at least one super-population as sampled by 1KG. This basis was further reduced, for each individual cancer-derived molecular profile, to genotypes at positions with high sequence coverage by high-quality reads in the profile. We then computed singular-value decomposition of the reduced 1KG genotype matrix and projected the genotype of the cancer-derived profile onto the first *D* of the resulting principal components. The ancestry for the profile was determined as that of the majority among the nearest *K* 1KG neighbors of the profile in this *D*-dimensional space [8]. For a subset of patients in each cohort we individually assessed the performance of the ancestry inference, as a function of the parameters *D* and *K*. This assessment was based, for each patient in the subset, on a large number of synthetic cancer-derived molecular profiles, as outlined in the Introduction, schematically described in ***Figure 4*** and explained in greater detail in the Methods section. The result was quantified, for a given *D, K* pair of parameters, as the area under receiver operating characteristic (AUROC) [31–33]. Both super-population-specific and overall AUROC values were computed in a range of *D, K* pairs, as illustrated in ***Figure 5*** for 10 PDAC patients and AMR-specific AUROC (the similar figures for all the cohorts and super-populations are in Figure S1). Optimal *D, K* pairs maximizing the overall AUROC were chosen. From this subset of patients we observed, for each cancer type considered and for each of the three molecular profiling modalities, an optimal range of *D* and *K* parameters where the performance of inference was consistently high in the subset and only weakly dependent on these parameters (Figure S1). We then selected and used, for the remainder of the patients with this cancer type and for this profiling modality, a pair *D* and *K* values from within the optimal range. As an additional validation of our parameter optimization procedure, we applied it to a set of cancer-free WES profiles of TCGA-OV patients. Comparing the resulting ancestry calls to the consensus calls (C5) by TCGA [7], we find the two to be in excellent agreement Table S3 and S4.

We also assessed the cohort-wide performance of our ancestry calls from original cancer-derived molecular data, by comparison to the gold standard of ancestry as determined from the matching cancer-free genotypes. For Beat AML and TCGA-OV patients, we performed ancestry inference from cancer-free patient exomes, using the same methodology as as we did for the cancer-derived sequences of these patients. In the case of PDAC, cancer-free whole-genome sequencing data were available, and used for the same purpose for a portion of the patient cohort. For all three cohorts, we summarize our cohort-wide findings in ***Table 1*** (we include similar tables for the synthetic data Tables S5-S7). Ancestry calls from both microarray- and exome-derived genotypes were recently published by TCGA consortium [7], and we also used these so-called consensus (C5 in the following) calls in our performance assessment for TCGA-OV (Table S3 and S4).

**Table 1.** Cohort-wide performance measures for super-population calls from cancer-derived molecular data, as compared to the matching cancer-free WES or (in the case of PDAC) WGS. A reliable estimate of the confidence intervals (CI) was not possible in the case of PDAC, due to the small number of cases with matching cancer-free genotypes.

| Study | D | K | Accuracy | 95% CI | AUROC | 95% CI |
| --- | --- | --- | --- | --- | --- | --- |
| TCGA-OV WES | 5 | 13 | 0.998 | 0.994-1 | 0.993 | 0.992-0.994 |
| TCGA-OV Panel | 4 | 12 | 0.984 | 0.972-0.996 | 0.966 | 0.965-0.967 |
| TCGA-OV RNA-seq | 7 | 12 | 0.993 | 0.983-1 | 0.977 | 0.975-0.979 |
| BeatAML WES | 5 | 13 | 0.989 | 0.978-1 | 0.978 | 0.976-0.980 |
| BeatAML Panel | 4 | 13 | 0.991 | 0.981-1 | 0.999 | 0.999-0.999 |
| BeatAML RNA-seq | 4 | 13 | 0.992 | 0.981-1 | 0.999 | 0.999-0.999 |
| PDAC WES | 8 | 13 | 1 | NA | NA | NA |
| PDAC Panel | 6 | 5 | 0.952 | 0.861-1 | 0.958 | NA |
| PDAC RNA-seq | 4 | 13 | 1 | NA | NA | NA |

We note that in the three patient cohorts we analyze here the sampling of patients with non-European ancestries is statistically insufficient for a purely cohort-based assessment of performance (***Table 2*** and Table S8). We therefore report cohort-wide overall but not super-population specific AUROC values. Using data synthesis, we are able to compensate for this data shortfall in non-European ancestries and estimate super-population specific AUROC, as explained above (Tables S9-S11 and Figure S1).

**Table 2.**
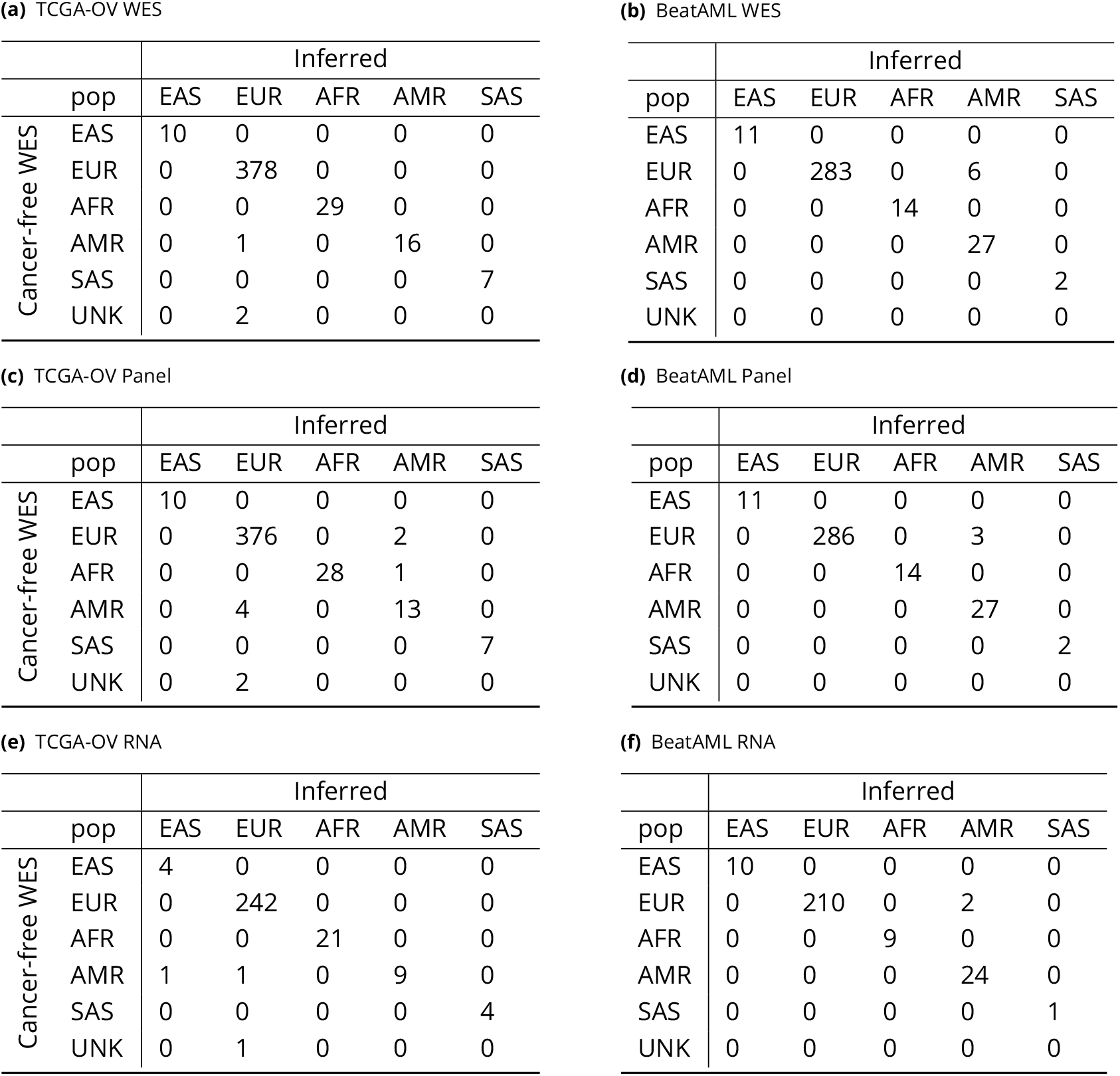
Confusion matrices comparing TCGA-OV or Beat AML patients’ super-population calls from the cancer-derived molecular profiles for the three profiling modalities (rows) to those from the matching cancer-free WES.

|  |  | Inferred |  |  |  |  |
| --- | --- | --- | --- | --- | --- | --- |
|  | pop | EAS | EUR | AFR | AMR | SAS |
| Cancer-free WES | EAS | 10 | 0 | 0 | 0 | 0 |
|  | EUR | 0 | 378 | 0 | 0 | 0 |
|  | AFR | 0 | 0 | 29 | 0 | 0 |
|  | AMR | 0 | 1 | 0 | 16 | 0 |
|  | SAS | 0 | 0 | 0 | 0 | 7 |
|  | UNK | 0 | 2 | 0 | 0 | 0 |

**(b)** BeatAML WES
|  |  | Inferred |  |  |  |  |
| --- | --- | --- | --- | --- | --- | --- |
|  | pop | EAS | EUR | AFR | AMR | SAS |
| Cancer-free WES | EAS | 11 | 0 | 0 | 0 | 0 |
|  | EUR | 0 | 283 | 0 | 6 | 0 |
|  | AFR | 0 | 0 | 14 | 0 | 0 |
|  | AMR | 0 | 0 | 0 | 27 | 0 |
|  | SAS | 0 | 0 | 0 | 0 | 2 |
|  | UNK | 0 | 0 | 0 | 0 | 0 |

**(c)** TCGA-OV Panel
|  |  | Inferred |  |  |  |  |
| --- | --- | --- | --- | --- | --- | --- |
|  | pop | EAS | EUR | AFR | AMR | SAS |
| Cancer-free WES | EAS | 10 | 0 | 0 | 0 | 0 |
|  | EUR | 0 | 376 | 0 | 2 | 0 |
|  | AFR | 0 | 0 | 28 | 1 | 0 |
|  | AMR | 0 | 4 | 0 | 13 | 0 |
|  | SAS | 0 | 0 | 0 | 0 | 7 |
|  | UNK | 0 | 2 | 0 | 0 | 0 |

**(d)** BeatAML Panel
|  |  | Inferred |  |  |  |  |
| --- | --- | --- | --- | --- | --- | --- |
|  | pop | EAS | EUR | AFR | AMR | SAS |
| Cancer-free WES | EAS | 11 | 0 | 0 | 0 | 0 |
|  | EUR | 0 | 286 | 0 | 3 | 0 |
|  | AFR | 0 | 0 | 14 | 0 | 0 |
|  | AMR | 0 | 0 | 0 | 27 | 0 |
|  | SAS | 0 | 0 | 0 | 0 | 2 |
|  | UNK | 0 | 0 | 0 | 0 | 0 |

**(e)** TCGA-OV RNA
|  |  | Inferred |  |  |  |  |
| --- | --- | --- | --- | --- | --- | --- |
|  | pop | EAS | EUR | AFR | AMR | SAS |
| Cancer-free WES | EAS | 4 | 0 | 0 | 0 | 0 |
|  | EUR | 0 | 242 | 0 | 0 | 0 |
|  | AFR | 0 | 0 | 21 | 0 | 0 |
|  | AMR | 1 | 1 | 0 | 9 | 0 |
|  | SAS | 0 | 0 | 0 | 0 | 4 |
|  | UNK | 0 | 1 | 0 | 0 | 0 |

|  |  | Inferred |  |  |  |  |
| --- | --- | --- | --- | --- | --- | --- |
|  | pop | EAS | EUR | AFR | AMR | SAS |
| Cancer-free WES | EAS | 10 | 0 | 0 | 0 | 0 |
|  | EUR | 0 | 210 | 0 | 2 | 0 |
|  | AFR | 0 | 0 | 9 | 0 | 0 |
|  | AMR | 0 | 0 | 0 | 24 | 0 |
|  | SAS | 0 | 0 | 0 | 0 | 1 |
|  | UNK | 0 | 0 | 0 | 0 | 0 |

The results of our analysis as presented in Tables S9-S11, lead to the following key observations. First, we demonstrate a consistently high performance of our inference procedure across all cohorts and profiling modalities. Second, the super-population specific performance was the highest for the European and both Asian super populations. The slightly lower accuracy as observed for the African and mixed American super-populations is likely due to a greater genetic variability within the African super-population and to a higher degree of (the predominantly European) admixture in both super-populations. Third, the optimal choice of the *D, K* inference parameters, in general, depends on an individual cancer-derived molecular profile, even within the same cancer type and profiling modality (Figure S1 B,G,L).

In order to examine whether our inference procedure is robust against variation in the sequence target coverage, we re-computed the ancestry calls for a subset of ten OV patients, with the cancer-derived whole-exome and RNA sequences of these patients down-sampled to between 75% and 10% of the original coverage. The results, presented in (Figure S2) exhibit no substantial sensitivity of the inference accuracy to the depth of coverage in this range.

## Discussion

With this work, we introduce a systematic approach to ancestry inference from cancer-derived molecular data. The approach is rooted in a combination of an established, extensively used PCA-based technique of ancestry inference with a central idea of inference parameter optimization using data synthesized *in silico*. Crucially, this combination permits a statistically rigorous assessment of inference accuracy for an individual cancer-derived molecular profile, with its unique biological (e.g. cancer type) and technical (e.g., sequencing depth and quality) properties. Synthetic data here are used as a substitute for a real-world set of molecular profiles sharing these properties and with known ground-truth genetic ancestry. It is unrealistic to expect such a real-world set to be available in all cases. Our tests of the resulting computational methodology on a representative subset of cancer-derived data demonstrate its accurate and robust performance. As we describe in detail in the Methods section, our data synthesis method relies on heuristic components for an estimate of the allele fractions throughout the cancer-derived profile. This estimate can be made more rigorous by using haplotypes in future implementations of the method, but the present version produces allele fractions in good agreement with published allele fractions (ASCAT2 results in [35, 36]).

A line of research and development initiated with this work must be extended in several directions. First, the performance of the methods presented must be examined more comprehensively across cancer types, and sequence properties, such as quality and depth. This task is computing-intensive but feasible given extensive, well annotated repositories of cancer-derived data, such as those resulting from TCGA Research Network [43] and ICGC [44] projects. For these, the genetic ancestry of the patients either is known or can be readily established using matching cancer-free molecular data. Second, an extension of our approach to additional profiling modalities should be examined. Chief among these are low-coverage whole-genome sequences commonly used for copy-number analysis and single-molecule, long-read sequences. Each of these presents unique challenges and opportunities for the ancestry inference: in the former, the sparsity of coverage is compensated by its whole-genome breadth; in the latter, the trade-off is between the high sequence error rate and the long-distance phasing afforded by long reads. Third, while the present work relied on PCA followed by nearest-neighbor classification for ancestry assessment, alternatives including UMAP for the former and Random Forest or Support Vector Machine for the latter exist and should be evaluated. Third, future method development should be extended beyond inference of global ancestry to that of local ancestry and ancestral admixture. Such an extension is particularly important in the study of cancer in strongly admixed populations, such as African and Latin Americans and may require more extensive reference data, in addition to the 1KG reference used here. Finally, beyond cancer, our methodology can be applied to inference from genomic data originating in any kind of fragmentary or damaged nucleic-acid specimens, such as those encountered in forensic, archaeological or paleontological contexts.

We anticipate the computational approach described here to have a major, two-fold, impact on investigation of links between ancestry and cancer. First, it will become possible to massively boost the statistical power of such studies by leveraging existing tumor-derived molecular data sets without matching germline sequences or ancestry annotation. Our search of the Gene Expression Omnibus (GEO) database alone has identified over 1,250 such data sets, containing RNA expression data for nearly 48,000 cancer tissue specimens. Such resources dwarf those of fully annotated repositories, such as TCGA and International Cancer Genome Consortium (ICGC) [44]. Other molecular data repositories are likely to contain resources of this category on a similar order of magnitude. Second, hundreds of thousands of tumor tissue specimens stored at multiple clinical centers constitute another major resource for ancestry-aware molecular studies of cancer. Here again, matching normal tissue specimens are often absent, and so is ethnic or racial annotation for the patients. According to a recent estimate [45] such annotation is missing in electronic health records of over 50% of patients. Inferential tools presented here will make these massive resources of archival tissues available for ancestry-oriented cancer research.

Multiple directions of exploratory and correlative analysis are open to pursuit with the accurate ancestry annotation made possible by the methods described here, even in the absence of matching cancer-free molecular data. Single-nucleotide and other small-scale somatic alterations may be identified in cancer-only exomes, both whole and restricted to specialized gene panels, using methods developed for this purpose [46] alongside databases of frequent somatic variants in cancer [18] and of frequent germline variants like gnomAD [47] and 1KG [42]. Copy number variants and losses of heterozygosity in cancer exomes are overwhelmingly somatic and may be determined computationally [48, 49]. Cancer RNA expression quantification is feasible in the absence of the germline genotype of the patient, including alllele- and isoform-specific analysis. These and similar genomic and transcriptional properties may be explored for associations with ancestral background of the patients.

## Acknowledgments

DAT is a distinguished scholar of the Lustgarten Foundation and Director of the Lustgarten Foundation-designated Laboratory of Pancreatic Cancer Research. DAT is also supported by the Cold Spring Harbor Laboratory Association, the New York Genome Center Polyethnic 1000 Project, the V Foundation (T2016-010), the Thompson Foundation, the Simons Foundation (552716), and the NIH (P30CA45508, P20CA192996, U10CA180944, U01CA224013, U01CA210240, R01CA188134, R01CA249002, and R01CA229699). AK’s work was supported by the New York Genome Center Polyethnic-1000 Major Grant, Simons Foundation award # 519054, Lustgarten Foundation OPT2 project award and by the Simons Center for Quantitative Biology at Cold Spring Harbor Laboratory. This work was performed with assistance from the US National Institutes of Health Grant S10OD028632-01. The results published here are in part based upon data generated by TCGA Research Network: https://www.cancer.gov/tcga. We thank Adam Siepel, Lloyd Trotman, Jeffrey Boyd, W. Richard Mc-Combie, Thomas Gingeras, Justin Kinney, Camila dos Santos, Michael Schatz, Louis Staudt, Michael Berger, David Solit and Samuel Aparicio for illuminating discussions.

