## Supplementary_Material for "Accurate and robust inference of genetic ancestry from cancer-derived molecular data across genomic platforms"

### Contents

**Public data used in the study** **3**

**Performance measures for ancestry inference** **4**

**Performance of genetic ancestry inference as a function of coverage** **28**

### List of Tables

1 Super-population and population representation in the 1KG reference data set, with no relatives among the individuals included. For the data synthesis, 30 genotypes were sampled from each 1KG population category, resulting in the sampling from each of the five super-populations as shown. . 3

2 Cancer-derived data used from the TCGA-OV, Beat AML and PDAC cohorts. . . . . 3

3 Cohort-wide performance measures for super-population inference from TCGA-OV molecular data, with the C5 (***Carrot-Zhang et al., 2020***) ancestry calls as the ground truth. . . . . 4

4 Confusion matrices for super-population calls from TCGA-OV cancer-derived data, in comparison to the C5 calls (***Carrot-Zhang et al., 2020***) . . . . . 5

5 Overall AUROC for 10 TCGA-OV patients, computed using data synthesis. . . . . 6

6 Overall AUROC for 10 Beat AML patients, computed using data synthesis. . . . . 7

7 Overall AUROC for 10 PDAC patients, computed using data synthesis. . . . . 8

8 Confusion matrices for super-population calls from PDAC cancer-derived data, in comparison to those from cancer-free WGS. . . . . 9

9 Super-population specific AUROC for 10 TCGA-OV patients, computed using data synthesis. . . . . 10

10 Super-population specific AUROC for 10 Beat AML patients, computed using data synthesis. . . . . . 11
11 Super-population specific AUROC for 10 PDAC patients, computed using data synthesis. . . . . . . 12

### Public data used in the study

| AFR | | AMR | | EAS | | EUR | | SAS | |
|---|---|---|---|---|---|---|---|---|---|
| pop | number | pop | number | pop | number | pop | number | pop | number |
| ACB | 95 | CLM | 94 | CDX | 96 | CEU | 96 | BEB | 85 |
| ASW | 50 | MXL | 62 | CHB | 106 | FIN | 105 | GIH | 103 |
| ESN | 96 | PEL | 84 | CHS | 101 | GBR | 96 | ITU | 101 |
| GWD | 109 | PUR | 104 | JPT | 105 | IBS | 107 | PJL | 91 |
| LWK | 88 | - | 0 | KHV | 99 | TSI | 108 | STU | 99 |
| MSL | 82 | - | 0 | - | 0 | - | 0 | - | 0 |
| YRI | 107 | - | 0 | - | 0 | - | 0 | - | 0 |
| Total | 627 | - | 344 | - | 507 | - | 512 | - | 479 |
| Sampled | 210 | - | 120 | - | 150 | - | 150 | - | 150 |

**Table 1.** Super-population and population representation in the 1KG reference data set, with no relatives among the individuals included. For the data synthesis, 30 genotypes were sampled from each 1KG population category, resulting in the sampling from each of the five super-populations as shown.

| Cohort | | Sample | |
|---|---|---|---|
| Cohort | Profiling modality | Source | Count |
| TCGA-OV | WES | Cancer | 453 |
| TCGA-OV | WES | Cancer-free | 450 |
| TCGA-OV | RNA-seq | Cancer | 376 |
| Beat AML | WES | Cancer | 343 |
| Beat AML | WES | Cancer-free | 532 |
| Beat AML | RNA-seq | Cancer | 430 |
| PDAC | WES | Cancer | 65 |
| PDAC | WGS | Cancer-free | 24 |
| PDAC | RNA-seq | Cancer | 40 |

**Table 2.** Cancer-derived data used from the TCGA-OV, Beat AML and PDAC cohorts.

### Performance measures for ancestry inference

| Source and modality | D | K | Accuracy | 95% CI | AUROC | 95% CI |
|---|---|---|---|---|---|---|
| Cancer-free WES | 5 | 13 | 0.986 | 0.975-0.997 | 0.994 | 0.994-0.995 |
| Cancer WES | 5 | 13 | 0.990 | 0.980-1 | 0.995 | 0.994-0.995 |
| Cancer panel | 4 | 12 | 0.991 | 0.982-1 | 0.995 | 0.994-0.995 |
| Cancer RNA-seq | 7 | 12 | 0.989 | 0.978-1 | 0.978 | 0.976-0.981 |

**Table 3.** Cohort-wide performance measures for super-population inference from TCGA-OV molecular data, with the C5 (***Carrot-Zhang et al., 2020***) ancestry calls as the ground truth.

**(a)** WES

| | | Inferred | | | | |
|---|---|---|---|---|---|---|
| | pop | EAS | EUR | AFR | AMR | SAS |
| C5 | EAS | 11 | 0 | 0 | 0 | 0 |
| | EUR | 0 | 386 | 0 | 4 | 0 |
| | AFR | 0 | 0 | 29 | 1 | 0 |
| | AMR | 0 | 0 | 0 | 1 | 0 |
| | SAS | 0 | 0 | 0 | 0 | 6 |

**(b)** RNA-seq

| | | Inferred | | | | |
|---|---|---|---|---|---|---|
| | pop | EAS | EUR | AFR | AMR | SAS |
| C5 | EAS | 7 | 1 | 0 | 0 | 0 |
| | EUR | 0 | 325 | 0 | 2 | 0 |
| | AFR | 0 | 0 | 23 | 1 | 0 |
| | AMR | 0 | 1 | 0 | 1 | 0 |
| | SAS | 0 | 0 | 0 | 0 | 4 |

**(c)** Panel

| | | Inferred | | | | |
|---|---|---|---|---|---|---|
| | pop | EAS | EUR | AFR | AMR | SAS |
| C5 | EAS | 11 | 0 | 0 | 0 | 0 |
| | EUR | 0 | 387 | 0 | 3 | 0 |
| | AFR | 0 | 0 | 29 | 1 | 0 |
| | AMR | 0 | 0 | 0 | 1 | 0 |
| | SAS | 0 | 0 | 0 | 0 | 6 |

**Table 4.** Confusion matrices for super-population calls from TCGA-OV cancer-derived data, in comparison to the C5 calls (***Carrot-Zhang et al., 2020***)

**Table 5.** Overall AUROC for 10 TCGA-OV patients, computed using data synthesis.

| | P | D | K | Accuracy | 95% CI | AUROC | 95% CI |
|---|---|---|---|---|---|---|---|
| WES | P1 | 5 | 13 | 0.9974359 | 0.994-1 | 0.9983631 | 0.998-0.998 |
| | P2 | 5 | 13 | 0.9974359 | 0.994-1 | 0.9983631 | 0.998-0.998 |
| | P3 | 5 | 13 | 0.9974359 | 0.994-1 | 0.9983631 | 0.998-0.998 |
| | P4 | 5 | 13 | 0.9974359 | 0.994-1 | 0.9983631 | 0.998-0.998 |
| | P5 | 5 | 13 | 0.9974359 | 0.994-1 | 0.9983631 | 0.998-0.998 |
| | P6 | 5 | 13 | 0.9974359 | 0.994-1 | 0.9983631 | 0.998-0.998 |
| | P7 | 5 | 13 | 0.9974359 | 0.994-1 | 0.9983631 | 0.998-0.998 |
| | P8 | 5 | 13 | 0.9974359 | 0.994-1 | 0.9983631 | 0.998-0.998 |
| | P9 | 5 | 13 | 0.9974359 | 0.994-1 | 0.9983631 | 0.998-0.998 |
| | P10 | 5 | 13 | 0.9974359 | 0.994-1 | 0.9983631 | 0.998-0.998 |
| Panel | P1 | 4 | 12 | 0.9923077 | 0.986-0.998 | 0.9944048 | 0.994-0.995 |
| | P2 | 4 | 12 | 0.9923077 | 0.986-0.998 | 0.9944048 | 0.994-0.995 |
| | P3 | 4 | 12 | 0.9923077 | 0.986-0.998 | 0.9944048 | 0.994-0.995 |
| | P4 | 4 | 12 | 0.9935897 | 0.988-0.999 | 0.9954464 | 0.995-0.996 |
| | P5 | 4 | 12 | 0.9935897 | 0.988-0.999 | 0.9952381 | 0.995-0.995 |
| | P6 | 4 | 12 | 0.9923077 | 0.986-0.998 | 0.9944048 | 0.994-0.995 |
| | P7 | 4 | 12 | 0.9948718 | 0.99-1 | 0.9962798 | 0.996-0.996 |
| | P8 | 4 | 12 | 0.9935897 | 0.988-0.999 | 0.9952381 | 0.995-0.995 |
| | P9 | 4 | 12 | 0.9910256 | 0.984-0.998 | 0.9935714 | 0.993-0.994 |
| | P10 | 4 | 12 | 0.9923077 | 0.986-0.998 | 0.9944048 | 0.994-0.995 |
| RNA-seq | P1 | 7 | 12 | 0.9974359 | 0.994-1 | 0.9983631 | 0.998-0.998 |
| | P2 | 7 | 12 | 0.9974359 | 0.994-1 | 0.9983631 | 0.998-0.998 |
| | P3 | 7 | 12 | 0.9974359 | 0.994-1 | 0.9983631 | 0.998-0.998 |
| | P4 | 7 | 12 | 0.9974359 | 0.994-1 | 0.9983631 | 0.998-0.998 |
| | P5 | 7 | 12 | 0.9974359 | 0.994-1 | 0.9983631 | 0.998-0.998 |
| | P6 | 7 | 12 | 0.9974359 | 0.994-1 | 0.9983631 | 0.998-0.998 |
| | P7 | 7 | 12 | 0.9974359 | 0.994-1 | 0.9983631 | 0.998-0.998 |
| | P8 | 7 | 12 | 0.9974359 | 0.994-1 | 0.9983631 | 0.998-0.998 |
| | P9 | 7 | 12 | 0.9974359 | 0.994-1 | 0.9983631 | 0.998-0.998 |
| | P10 | 7 | 12 | 0.9974359 | 0.994-1 | 0.9983631 | 0.998-0.998 |

**Table 6.** Overall AUROC for 10 Beat AML patients, computed using data synthesis.

| | P | D | K | Accuracy | 95% CI | AUROC | 95% CI |
|---|---|---|---|---|---|---|---|
| WES | P1 | 5 | 13 | 0.9974359 | 0.994-1 | 0.9983631 | 0.998-0.998 |
| | P2 | 5 | 13 | 0.9974359 | 0.994-1 | 0.9983631 | 0.998-0.998 |
| | P3 | 5 | 13 | 0.9974359 | 0.994-1 | 0.9983631 | 0.998-0.998 |
| | P4 | 5 | 13 | 0.9974359 | 0.994-1 | 0.9983631 | 0.998-0.998 |
| | P5 | 5 | 13 | 0.9974359 | 0.994-1 | 0.9983631 | 0.998-0.998 |
| | P6 | 5 | 13 | 0.9974359 | 0.994-1 | 0.9983631 | 0.998-0.998 |
| | P7 | 5 | 13 | 0.9974359 | 0.994-1 | 0.9983631 | 0.998-0.998 |
| | P8 | 5 | 13 | 0.9974359 | 0.994-1 | 0.9983631 | 0.998-0.998 |
| | P9 | 5 | 13 | 0.9974359 | 0.994-1 | 0.9983631 | 0.998-0.998 |
| | P10 | 5 | 13 | 0.9974359 | 0.994-1 | 0.9983631 | 0.998-0.998 |
| Panel | P1 | 4 | 13 | 0.9961538 | 0.992-1 | 0.9973214 | 0.997-0.997 |
| | P2 | 4 | 13 | 0.9935897 | 0.988-0.999 | 0.9954464 | 0.995-0.996 |
| | P3 | 4 | 13 | 0.9961538 | 0.992-1 | 0.9973214 | 0.997-0.997 |
| | P4 | 4 | 13 | 0.9961538 | 0.992-1 | 0.9973214 | 0.997-0.997 |
| | P5 | 4 | 13 | 0.9961538 | 0.992-1 | 0.9973214 | 0.997-0.997 |
| | P6 | 4 | 13 | 0.9935897 | 0.988-0.999 | 0.9956845 | 0.996-0.996 |
| | P7 | 4 | 13 | 0.9948718 | 0.99-1 | 0.9962798 | 0.996-0.996 |
| | P8 | 4 | 13 | 0.9948718 | 0.99-1 | 0.9967262 | 0.997-0.997 |
| | P9 | 4 | 13 | 0.9935897 | 0.988-0.999 | 0.9956845 | 0.996-0.996 |
| | P10 | 4 | 13 | 0.9935897 | 0.988-0.999 | 0.9956845 | 0.996-0.996 |
| RNA-seq | P1 | 4 | 13 | 0.9974359 | 0.994-1 | 0.9983631 | 0.998-0.998 |
| | P2 | 4 | 13 | 0.9974359 | 0.994-1 | 0.9983631 | 0.998-0.998 |
| | P3 | 4 | 13 | 0.9974359 | 0.994-1 | 0.9983631 | 0.998-0.998 |
| | P4 | 4 | 13 | 0.9974359 | 0.994-1 | 0.9983631 | 0.998-0.998 |
| | P5 | 4 | 13 | 0.9974359 | 0.994-1 | 0.9983631 | 0.998-0.998 |
| | P6 | 4 | 13 | 0.9974359 | 0.994-1 | 0.9983631 | 0.998-0.998 |
| | P7 | 4 | 13 | 0.9974359 | 0.994-1 | 0.9983631 | 0.998-0.998 |
| | P8 | 4 | 13 | 0.9974359 | 0.994-1 | 0.9983631 | 0.998-0.998 |
| | P9 | 4 | 13 | 0.9974359 | 0.994-1 | 0.9983631 | 0.998-0.998 |
| | P10 | 4 | 13 | 0.9974359 | 0.994-1 | 0.9983631 | 0.998-0.998 |

**Table 7.** Overall AUROC for 10 PDAC patients, computed using data synthesis.

| | P | D | K | Accuracy | 95% CI | AUROC | 95% CI |
|---|---|---|---|---|---|---|---|
| WES | P1 | 8 | 13 | 0.9974359 | 0.994-1 | 0.9983631 | 0.998-0.998 |
| | P2 | 8 | 13 | 0.9974359 | 0.994-1 | 0.9983631 | 0.998-0.998 |
| | P3 | 8 | 13 | 0.9974359 | 0.994-1 | 0.9983631 | 0.998-0.998 |
| | P4 | 8 | 13 | 0.9974359 | 0.994-1 | 0.9983631 | 0.998-0.998 |
| | P5 | 8 | 13 | 0.9974359 | 0.994-1 | 0.9983631 | 0.998-0.998 |
| | P6 | 8 | 13 | 0.9974359 | 0.994-1 | 0.9983631 | 0.998-0.998 |
| | P7 | 8 | 13 | 0.9974359 | 0.994-1 | 0.9983631 | 0.998-0.998 |
| | P8 | 8 | 13 | 0.9974359 | 0.994-1 | 0.9983631 | 0.998-0.998 |
| | P9 | 8 | 13 | 0.9974359 | 0.994-1 | 0.9983631 | 0.998-0.998 |
| | P10 | 8 | 13 | 0.9974359 | 0.994-1 | 0.9983631 | 0.998-0.998 |
| Panel | P1 | 6 | 5 | 0.9923077 | 0.986-0.998 | 0.9950595 | 0.995-0.995 |
| | P2 | 6 | 5 | 0.9948718 | 0.99-1 | 0.9964881 | 0.996-0.997 |
| | P3 | 6 | 5 | 0.9935897 | 0.988-0.999 | 0.9958929 | 0.996-0.996 |
| | P4 | 6 | 5 | 0.9948718 | 0.99-1 | 0.9964881 | 0.996-0.997 |
| | P5 | 6 | 5 | 0.9961538 | 0.992-1 | 0.9973214 | 0.997-0.997 |
| | P6 | 6 | 5 | 0.9923077 | 0.986-0.998 | 0.9950595 | 0.995-0.995 |
| | P7 | 6 | 5 | 0.9961538 | 0.992-1 | 0.9973214 | 0.997-0.997 |
| | P8 | 6 | 5 | 0.9935897 | 0.988-0.999 | 0.9958929 | 0.996-0.996 |
| | P9 | 6 | 5 | 0.9961538 | 0.992-1 | 0.9975298 | 0.997-0.998 |
| | P10 | 6 | 5 | 0.9948718 | 0.99-1 | 0.9964881 | 0.996-0.997 |
| RNA-seq | P1 | 5 | 11 | 0.9974359 | 0.994-1 | 0.9983631 | 0.998-0.998 |
| | P2 | 5 | 11 | 0.9974359 | 0.994-1 | 0.9983631 | 0.998-0.998 |
| | P3 | 5 | 11 | 0.9974359 | 0.994-1 | 0.9983631 | 0.998-0.998 |
| | P4 | 5 | 11 | 0.9974359 | 0.994-1 | 0.9983631 | 0.998-0.998 |
| | P5 | 5 | 11 | 0.9974359 | 0.994-1 | 0.9983631 | 0.998-0.998 |
| | P6 | 5 | 11 | 0.9974359 | 0.994-1 | 0.9983631 | 0.998-0.998 |
| | P7 | 5 | 11 | 0.9974359 | 0.994-1 | 0.9983631 | 0.998-0.998 |
| | P8 | 5 | 11 | 0.9974359 | 0.994-1 | 0.9983631 | 0.998-0.998 |
| | P9 | 5 | 11 | 0.9974359 | 0.994-1 | 0.9983631 | 0.998-0.998 |
| | P10 | 5 | 11 | 0.9974359 | 0.994-1 | 0.9983631 | 0.998-0.998 |

**(a)** WES

| | | Inferred | | | | |
|---|---|---|---|---|---|---|
| | pop | EAS | EUR | AFR | AMR | SAS |
| Cancer-free WGS | EAS | 1 | 0 | 0 | 0 | 0 |
| | EUR | 0 | 15 | 0 | 0 | 0 |
| | AFR | 0 | 0 | 2 | 0 | 0 |
| | AMR | 0 | 0 | 0 | 2 | 0 |
| | SAS | 0 | 0 | 0 | 0 | 1 |

**(b)** Panel

| | | Inferred | | | | |
|---|---|---|---|---|---|---|
| | pop | EAS | EUR | AFR | AMR | SAS |
| Cancer-free WGS | EAS | 1 | 0 | 0 | 0 | 0 |
| | EUR | 0 | 15 | 0 | 0 | 0 |
| | AFR | 0 | 0 | 2 | 0 | 0 |
| | AMR | 0 | 1 | 0 | 1 | 0 |
| | SAS | 0 | 0 | 0 | 0 | 1 |

**(c)** RNA-seq

| | | Inferred | | | | |
|---|---|---|---|---|---|---|
| | pop | EAS | EUR | AFR | AMR | SAS |
| Cancer-free WGS | EAS | 1 | 0 | 0 | 0 | 0 |
| | EUR | 0 | 13 | 0 | 0 | 0 |
| | AFR | 0 | 0 | 2 | 0 | 0 |
| | AMR | 0 | 0 | 0 | 2 | 0 |
| | SAS | 0 | 0 | 0 | 0 | 1 |

**Table 8.** Confusion matrices for super-population calls from PDAC cancer-derived data, in comparison to those from cancer-free WGS.

**Table 9.** Super-population specific AUROC for 10 TCGA-OV patients, computed using data synthesis.

| | P | EAS | 95% CI | EUR | 95% CI | AFR | 95% CI | AMR | 95% CI | SAS | 95% CI |
|---|---|---|---|---|---|---|---|---|---|---|---|
| WES | P1 | 1 | 1-1 | 1.000 | 1-1 | 0.997 | 0.992-1 | 0.995 | 0.987-1 | 1 | 1-1 |
| | P2 | 1 | 1-1 | 1.000 | 1-1 | 0.997 | 0.992-1 | 0.995 | 0.987-1 | 1 | 1-1 |
| | P3 | 1 | 1-1 | 1.000 | 1-1 | 0.997 | 0.992-1 | 0.995 | 0.987-1 | 1 | 1-1 |
| | P4 | 1 | 1-1 | 1.000 | 1-1 | 0.997 | 0.992-1 | 0.995 | 0.987-1 | 1 | 1-1 |
| | P5 | 1 | 1-1 | 1.000 | 1-1 | 0.997 | 0.992-1 | 0.995 | 0.987-1 | 1 | 1-1 |
| | P6 | 1 | 1-1 | 1.000 | 1-1 | 0.997 | 0.992-1 | 0.995 | 0.987-1 | 1 | 1-1 |
| | P7 | 1 | 1-1 | 1.000 | 1-1 | 0.997 | 0.992-1 | 0.995 | 0.987-1 | 1 | 1-1 |
| | P8 | 1 | 1-1 | 1.000 | 1-1 | 0.997 | 0.992-1 | 0.995 | 0.987-1 | 1 | 1-1 |
| | P9 | 1 | 1-1 | 1.000 | 1-1 | 0.997 | 0.992-1 | 0.995 | 0.987-1 | 1 | 1-1 |
| | P10 | 1 | 1-1 | 1.000 | 1-1 | 0.997 | 0.992-1 | 0.995 | 0.987-1 | 1 | 1-1 |
| Panel | P1 | 1 | 1-1 | 0.994 | 0.987-1 | 0.997 | 0.992-1 | 0.982 | 0.966-0.998 | 1 | 1-1 |
| | P2 | 1 | 1-1 | 0.994 | 0.987-1 | 0.997 | 0.992-1 | 0.982 | 0.966-0.998 | 1 | 1-1 |
| | P3 | 1 | 1-1 | 0.994 | 0.987-1 | 0.997 | 0.992-1 | 0.982 | 0.966-0.998 | 1 | 1-1 |
| | P4 | 1 | 1-1 | 0.995 | 0.988-1 | 0.997 | 0.992-1 | 0.986 | 0.972-1 | 1 | 1-1 |
| | P5 | 1 | 1-1 | 0.998 | 0.995-1 | 0.997 | 0.992-1 | 0.983 | 0.966-0.999 | 1 | 1-1 |
| | P6 | 1 | 1-1 | 0.994 | 0.987-1 | 0.997 | 0.992-1 | 0.982 | 0.966-0.998 | 1 | 1-1 |
| | P7 | 1 | 1-1 | 0.998 | 0.996-1 | 0.997 | 0.992-1 | 0.987 | 0.973-1 | 1 | 1-1 |
| | P8 | 1 | 1-1 | 0.998 | 0.995-1 | 0.997 | 0.992-1 | 0.983 | 0.966-0.999 | 1 | 1-1 |
| | P9 | 1 | 1-1 | 0.991 | 0.981-1 | 0.997 | 0.992-1 | 0.981 | 0.965-0.997 | 1 | 1-1 |
| | P10 | 1 | 1-1 | 0.994 | 0.987-1 | 0.997 | 0.992-1 | 0.982 | 0.966-0.998 | 1 | 1-1 |
| RNA-seq | P1 | 1 | 1-1 | 1.000 | 1-1 | 0.997 | 0.992-1 | 0.995 | 0.987-1 | 1 | 1-1 |
| | P2 | 1 | 1-1 | 1.000 | 1-1 | 0.997 | 0.992-1 | 0.995 | 0.987-1 | 1 | 1-1 |
| | P3 | 1 | 1-1 | 1.000 | 1-1 | 0.997 | 0.992-1 | 0.995 | 0.987-1 | 1 | 1-1 |
| | P4 | 1 | 1-1 | 1.000 | 1-1 | 0.997 | 0.992-1 | 0.995 | 0.987-1 | 1 | 1-1 |
| | P5 | 1 | 1-1 | 1.000 | 1-1 | 0.997 | 0.992-1 | 0.995 | 0.987-1 | 1 | 1-1 |
| | P6 | 1 | 1-1 | 1.000 | 1-1 | 0.997 | 0.992-1 | 0.995 | 0.987-1 | 1 | 1-1 |
| | P7 | 1 | 1-1 | 1.000 | 1-1 | 0.997 | 0.992-1 | 0.995 | 0.987-1 | 1 | 1-1 |
| | P8 | 1 | 1-1 | 1.000 | 1-1 | 0.997 | 0.992-1 | 0.995 | 0.987-1 | 1 | 1-1 |
| | P9 | 1 | 1-1 | 1.000 | 1-1 | 0.997 | 0.992-1 | 0.995 | 0.987-1 | 1 | 1-1 |
| | P10 | 1 | 1-1 | 1.000 | 1-1 | 0.997 | 0.992-1 | 0.995 | 0.987-1 | 1 | 1-1 |

**Table 10.** Super-population specific AUROC for 10 Beat AML patients, computed using data synthesis.

| | P | EAS | 95% CI | EUR | 95% CI | AFR | 95% CI | AMR | 95% CI | SAS | 95% CI |
|---|---|---|---|---|---|---|---|---|---|---|---|
| WES | P1 | 1 | 1-1 | 1.000 | 1-1 | 0.997 | 0.992-1 | 0.995 | 0.987-1 | 1 | 1-1 |
| | P2 | 1 | 1-1 | 1.000 | 1-1 | 0.997 | 0.992-1 | 0.995 | 0.987-1 | 1 | 1-1 |
| | P3 | 1 | 1-1 | 1.000 | 1-1 | 0.997 | 0.992-1 | 0.995 | 0.987-1 | 1 | 1-1 |
| | P4 | 1 | 1-1 | 1.000 | 1-1 | 0.997 | 0.992-1 | 0.995 | 0.987-1 | 1 | 1-1 |
| | P5 | 1 | 1-1 | 1.000 | 1-1 | 0.997 | 0.992-1 | 0.995 | 0.987-1 | 1 | 1-1 |
| | P6 | 1 | 1-1 | 1.000 | 1-1 | 0.997 | 0.992-1 | 0.995 | 0.987-1 | 1 | 1-1 |
| | P7 | 1 | 1-1 | 1.000 | 1-1 | 0.997 | 0.992-1 | 0.995 | 0.987-1 | 1 | 1-1 |
| | P8 | 1 | 1-1 | 1.000 | 1-1 | 0.997 | 0.992-1 | 0.995 | 0.987-1 | 1 | 1-1 |
| | P9 | 1 | 1-1 | 1.000 | 1-1 | 0.997 | 0.992-1 | 0.995 | 0.987-1 | 1 | 1-1 |
| | P10 | 1 | 1-1 | 1.000 | 1-1 | 0.997 | 0.992-1 | 0.995 | 0.987-1 | 1 | 1-1 |
| Panel | P1 | 1 | 1-1 | 0.999 | 0.998-1 | 0.997 | 0.992-1 | 0.991 | 0.979-1 | 1 | 1-1 |
| | P2 | 1 | 1-1 | 0.995 | 0.988-1 | 0.997 | 0.992-1 | 0.986 | 0.972-1 | 1 | 1-1 |
| | P3 | 1 | 1-1 | 0.999 | 0.998-1 | 0.997 | 0.992-1 | 0.991 | 0.979-1 | 1 | 1-1 |
| | P4 | 1 | 1-1 | 0.999 | 0.998-1 | 0.997 | 0.992-1 | 0.991 | 0.979-1 | 1 | 1-1 |
| | P5 | 1 | 1-1 | 0.999 | 0.998-1 | 0.997 | 0.992-1 | 0.991 | 0.979-1 | 1 | 1-1 |
| | P6 | 1 | 1-1 | 0.998 | 0.996-1 | 0.994 | 0.988-1 | 0.986 | 0.972-1 | 1 | 1-1 |
| | P7 | 1 | 1-1 | 0.998 | 0.996-1 | 0.997 | 0.992-1 | 0.987 | 0.973-1 | 1 | 1-1 |
| | P8 | 1 | 1-1 | 0.999 | 0.998-1 | 0.994 | 0.988-1 | 0.990 | 0.978-1 | 1 | 1-1 |
| | P9 | 1 | 1-1 | 0.998 | 0.996-1 | 0.994 | 0.988-1 | 0.986 | 0.972-1 | 1 | 1-1 |
| | P10 | 1 | 1-1 | 0.998 | 0.996-1 | 0.994 | 0.988-1 | 0.986 | 0.972-1 | 1 | 1-1 |
| RNA-seq | P1 | 1 | 1-1 | 1.000 | 1-1 | 0.997 | 0.992-1 | 0.995 | 0.987-1 | 1 | 1-1 |
| | P2 | 1 | 1-1 | 1.000 | 1-1 | 0.997 | 0.992-1 | 0.995 | 0.987-1 | 1 | 1-1 |
| | P3 | 1 | 1-1 | 1.000 | 1-1 | 0.997 | 0.992-1 | 0.995 | 0.987-1 | 1 | 1-1 |
| | P4 | 1 | 1-1 | 1.000 | 1-1 | 0.997 | 0.992-1 | 0.995 | 0.987-1 | 1 | 1-1 |
| | P5 | 1 | 1-1 | 1.000 | 1-1 | 0.997 | 0.992-1 | 0.995 | 0.987-1 | 1 | 1-1 |
| | P6 | 1 | 1-1 | 1.000 | 1-1 | 0.997 | 0.992-1 | 0.995 | 0.987-1 | 1 | 1-1 |
| | P7 | 1 | 1-1 | 1.000 | 1-1 | 0.997 | 0.992-1 | 0.995 | 0.987-1 | 1 | 1-1 |
| | P8 | 1 | 1-1 | 1.000 | 1-1 | 0.997 | 0.992-1 | 0.995 | 0.987-1 | 1 | 1-1 |
| | P9 | 1 | 1-1 | 1.000 | 1-1 | 0.997 | 0.992-1 | 0.995 | 0.987-1 | 1 | 1-1 |
| | P10 | 1 | 1-1 | 1.000 | 1-1 | 0.997 | 0.992-1 | 0.995 | 0.987-1 | 1 | 1-1 |

**Table 11.** Super-population specific AUROC for 10 PDAC patients, computed using data synthesis.

| | P | EAS | 95% CI | EUR | 95% CI | AFR | 95% CI | AMR | 95% CI | SAS | 95% CI |
|---|---|---|---|---|---|---|---|---|---|---|---|
| WES | P1 | 1 | 1-1 | 1.000 | 1-1 | 0.997 | 0.992-1 | 0.995 | 0.987-1 | 1 | 1-1 |
| | P2 | 1 | 1-1 | 1.000 | 1-1 | 0.997 | 0.992-1 | 0.995 | 0.987-1 | 1 | 1-1 |
| | P3 | 1 | 1-1 | 1.000 | 1-1 | 0.997 | 0.992-1 | 0.995 | 0.987-1 | 1 | 1-1 |
| | P4 | 1 | 1-1 | 1.000 | 1-1 | 0.997 | 0.992-1 | 0.995 | 0.987-1 | 1 | 1-1 |
| | P5 | 1 | 1-1 | 1.000 | 1-1 | 0.997 | 0.992-1 | 0.995 | 0.987-1 | 1 | 1-1 |
| | P6 | 1 | 1-1 | 1.000 | 1-1 | 0.997 | 0.992-1 | 0.995 | 0.987-1 | 1 | 1-1 |
| | P7 | 1 | 1-1 | 1.000 | 1-1 | 0.997 | 0.992-1 | 0.995 | 0.987-1 | 1 | 1-1 |
| | P8 | 1 | 1-1 | 1.000 | 1-1 | 0.997 | 0.992-1 | 0.995 | 0.987-1 | 1 | 1-1 |
| | P9 | 1 | 1-1 | 1.000 | 1-1 | 0.997 | 0.992-1 | 0.995 | 0.987-1 | 1 | 1-1 |
| | P10 | 1 | 1-1 | 1.000 | 1-1 | 0.997 | 0.992-1 | 0.995 | 0.987-1 | 1 | 1-1 |
| Panel | P1 | 1 | 1-1 | 0.993 | 0.983-1 | 0.994 | 0.988-1 | 0.989 | 0.977-1 | 1 | 1-1 |
| | P2 | 1 | 1-1 | 0.996 | 0.989-1 | 0.997 | 0.992-1 | 0.990 | 0.978-1 | 1 | 1-1 |
| | P3 | 1 | 1-1 | 0.996 | 0.989-1 | 0.994 | 0.988-1 | 0.989 | 0.978-1 | 1 | 1-1 |
| | P4 | 1 | 1-1 | 0.996 | 0.989-1 | 0.997 | 0.992-1 | 0.990 | 0.978-1 | 1 | 1-1 |
| | P5 | 1 | 1-1 | 0.999 | 0.998-1 | 0.997 | 0.992-1 | 0.991 | 0.979-1 | 1 | 1-1 |
| | P6 | 1 | 1-1 | 0.993 | 0.983-1 | 0.994 | 0.988-1 | 0.989 | 0.977-1 | 1 | 1-1 |
| | P7 | 1 | 1-1 | 0.999 | 0.998-1 | 0.997 | 0.992-1 | 0.991 | 0.979-1 | 1 | 1-1 |
| | P8 | 1 | 1-1 | 0.996 | 0.989-1 | 0.994 | 0.988-1 | 0.989 | 0.978-1 | 1 | 1-1 |
| | P9 | 1 | 1-1 | 0.997 | 0.99-1 | 0.997 | 0.992-1 | 0.994 | 0.986-1 | 1 | 1-1 |
| | P10 | 1 | 1-1 | 0.996 | 0.989-1 | 0.997 | 0.992-1 | 0.990 | 0.978-1 | 1 | 1-1 |
| RNA-seq | P1 | 1 | 1-1 | 1.000 | 1-1 | 0.997 | 0.992-1 | 0.995 | 0.987-1 | 1 | 1-1 |
| | P2 | 1 | 1-1 | 1.000 | 1-1 | 0.997 | 0.992-1 | 0.995 | 0.987-1 | 1 | 1-1 |
| | P3 | 1 | 1-1 | 1.000 | 1-1 | 0.997 | 0.992-1 | 0.995 | 0.987-1 | 1 | 1-1 |
| | P4 | 1 | 1-1 | 1.000 | 1-1 | 0.997 | 0.992-1 | 0.995 | 0.987-1 | 1 | 1-1 |
| | P5 | 1 | 1-1 | 1.000 | 1-1 | 0.997 | 0.992-1 | 0.995 | 0.987-1 | 1 | 1-1 |
| | P6 | 1 | 1-1 | 1.000 | 1-1 | 0.997 | 0.992-1 | 0.995 | 0.987-1 | 1 | 1-1 |
| | P7 | 1 | 1-1 | 1.000 | 1-1 | 0.997 | 0.992-1 | 0.995 | 0.987-1 | 1 | 1-1 |
| | P8 | 1 | 1-1 | 1.000 | 1-1 | 0.997 | 0.992-1 | 0.995 | 0.987-1 | 1 | 1-1 |
| | P9 | 1 | 1-1 | 1.000 | 1-1 | 0.997 | 0.992-1 | 0.995 | 0.987-1 | 1 | 1-1 |
| | P10 | 1 | 1-1 | 1.000 | 1-1 | 0.997 | 0.992-1 | 0.995 | 0.987-1 | 1 | 1-1 |

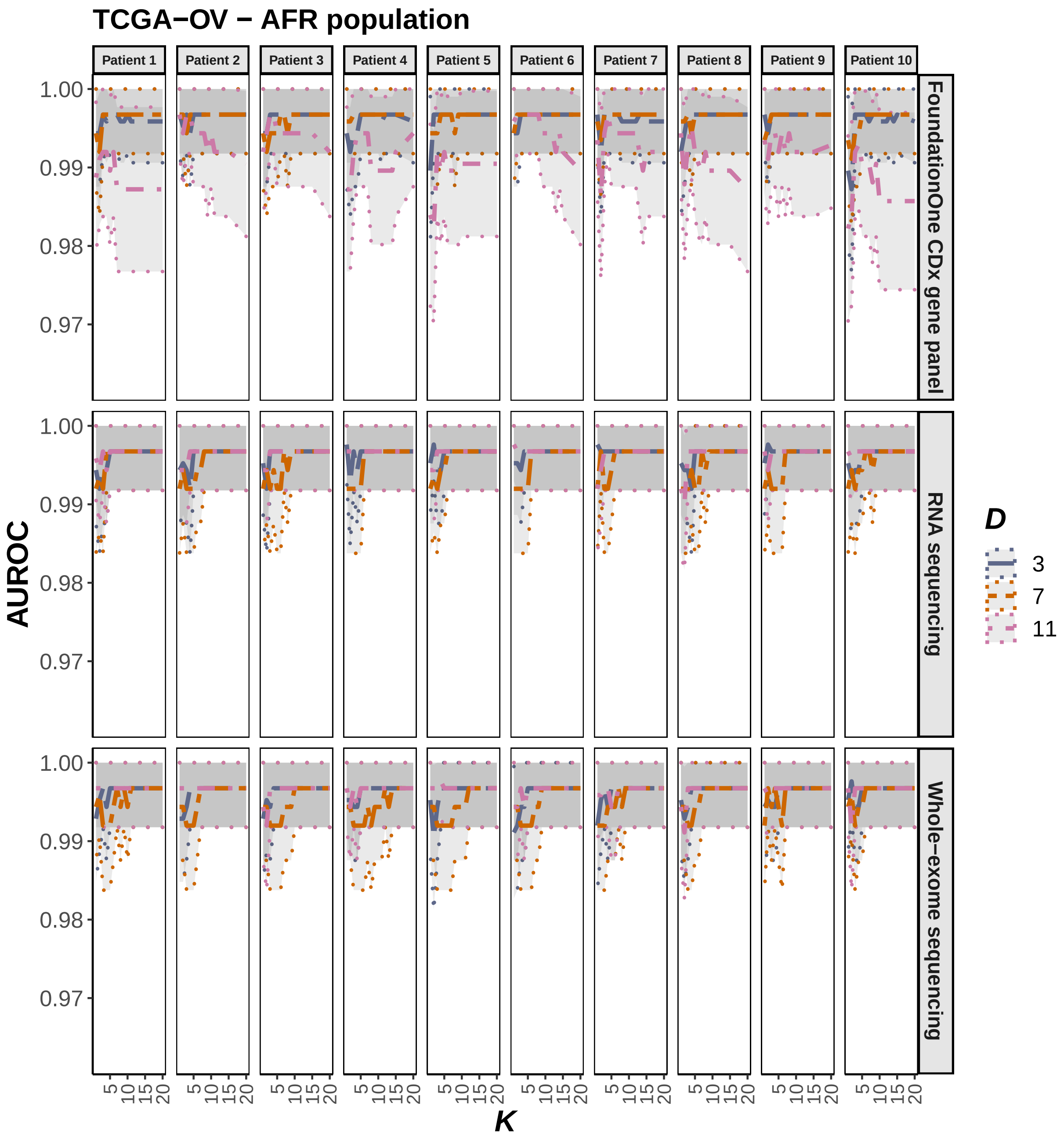

**Figure 1. A)** Dependence of AFR-specific AUROC on the inference parameters $D$ and $K$, computed using data synthesis for 10 TCGA-OV patients and the three profiling modalities: WES, RNA-seq and FoundationOne® CDx panels. The central AUROC values are shown in solid, and the 95% CI in dashed, lines.

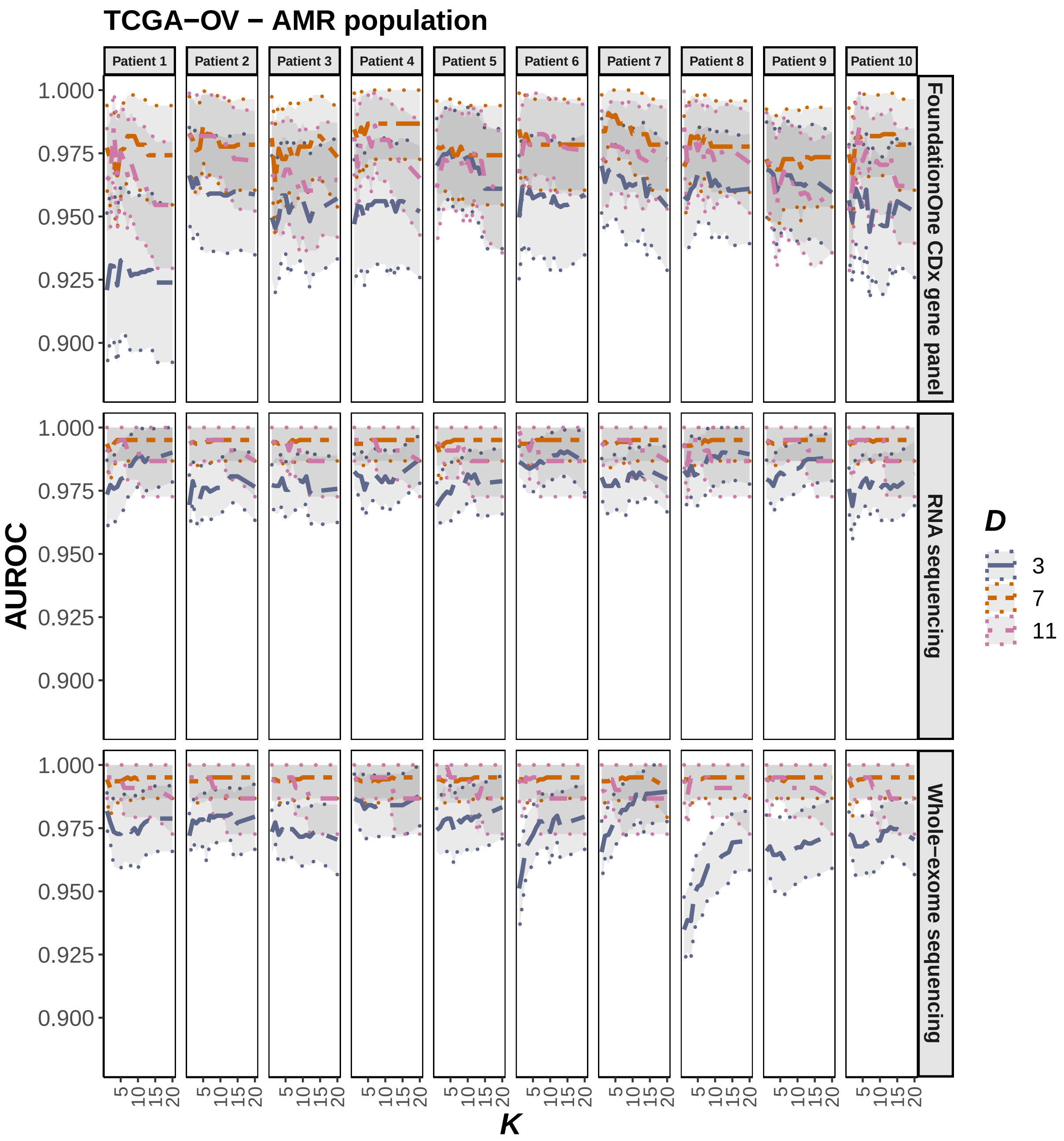

**Figure 1. B)** Dependence of AMR-specific AUROC on the inference parameters $D$ and $K$, computed using data synthesis for 10 TCGA-OV patients and the three profiling modalities: WES, RNA-seq and FoundationOne® CDx panels. The central AUROC values are shown in solid, and the 95% CI in dashed, lines.

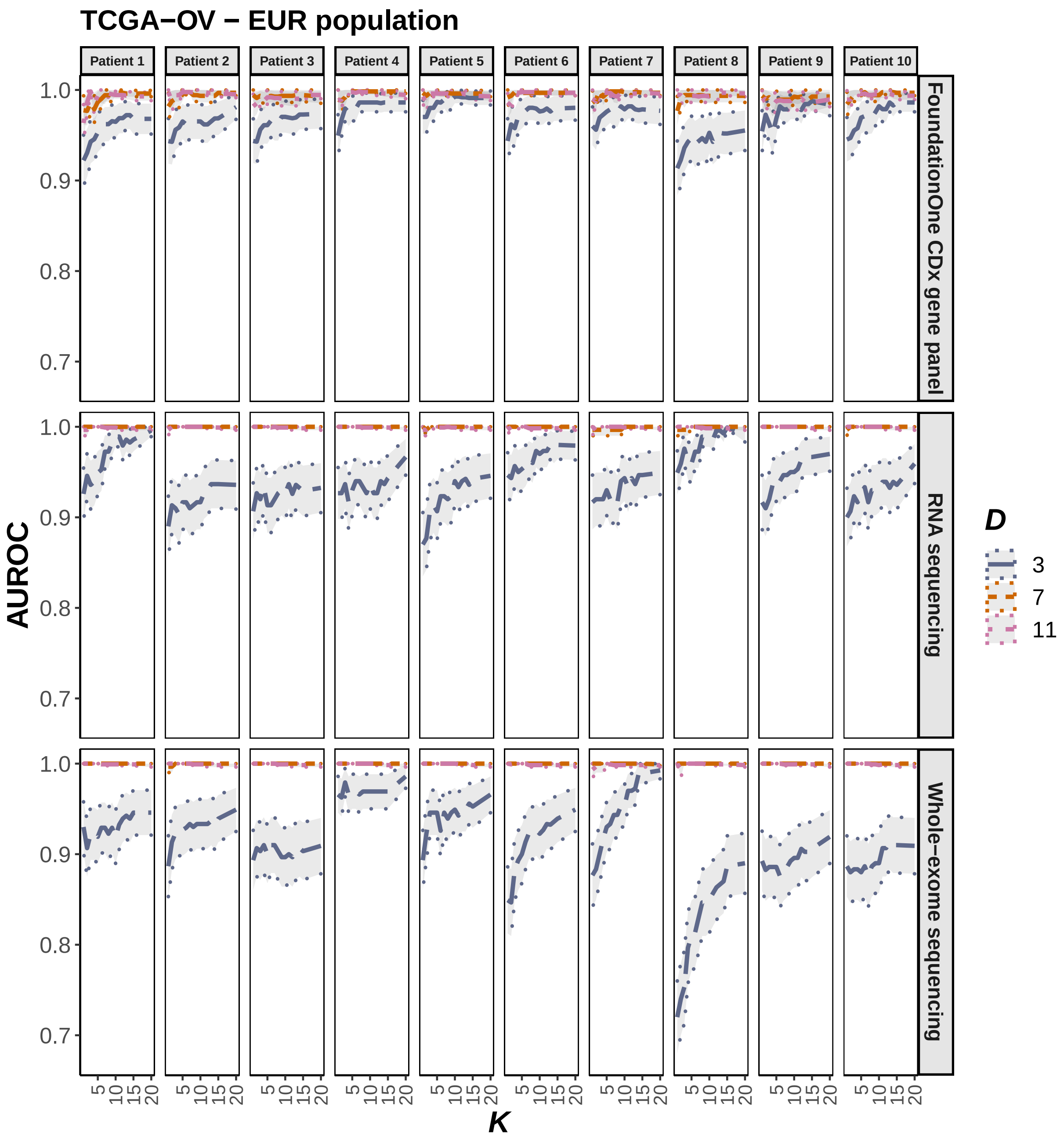

**Figure 1. C)** Dependence of EUR-specific AUROC on the inference parameters $D$ and $K$, computed using data synthesis for 10 TCGA-OV patients and the three profiling modalities: WES, RNA-seq and FoundationOne® CDx panels. The central AUROC values are shown in solid, and the 95% CI in dashed, lines.

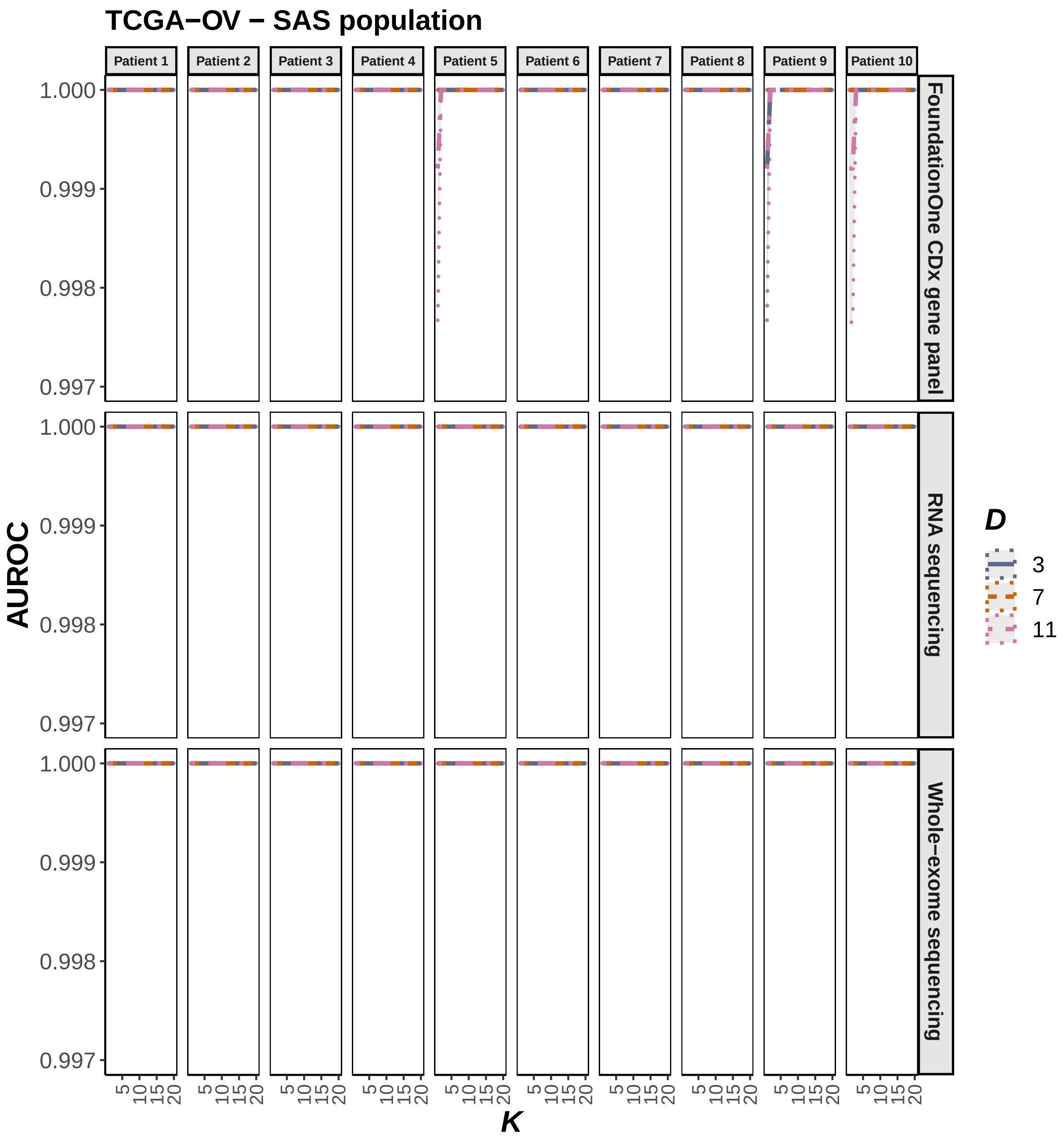

**Figure 1. D)** Dependence of SAS-specific AUROC on the inference parameters $D$ and $K$, computed using data synthesis for 10 TCGA-OV patients and the three profiling modalities: WES, RNA-seq and FoundationOne® CDx panels. The central AUROC values are shown in solid, and the 95% CI in dashed, lines.

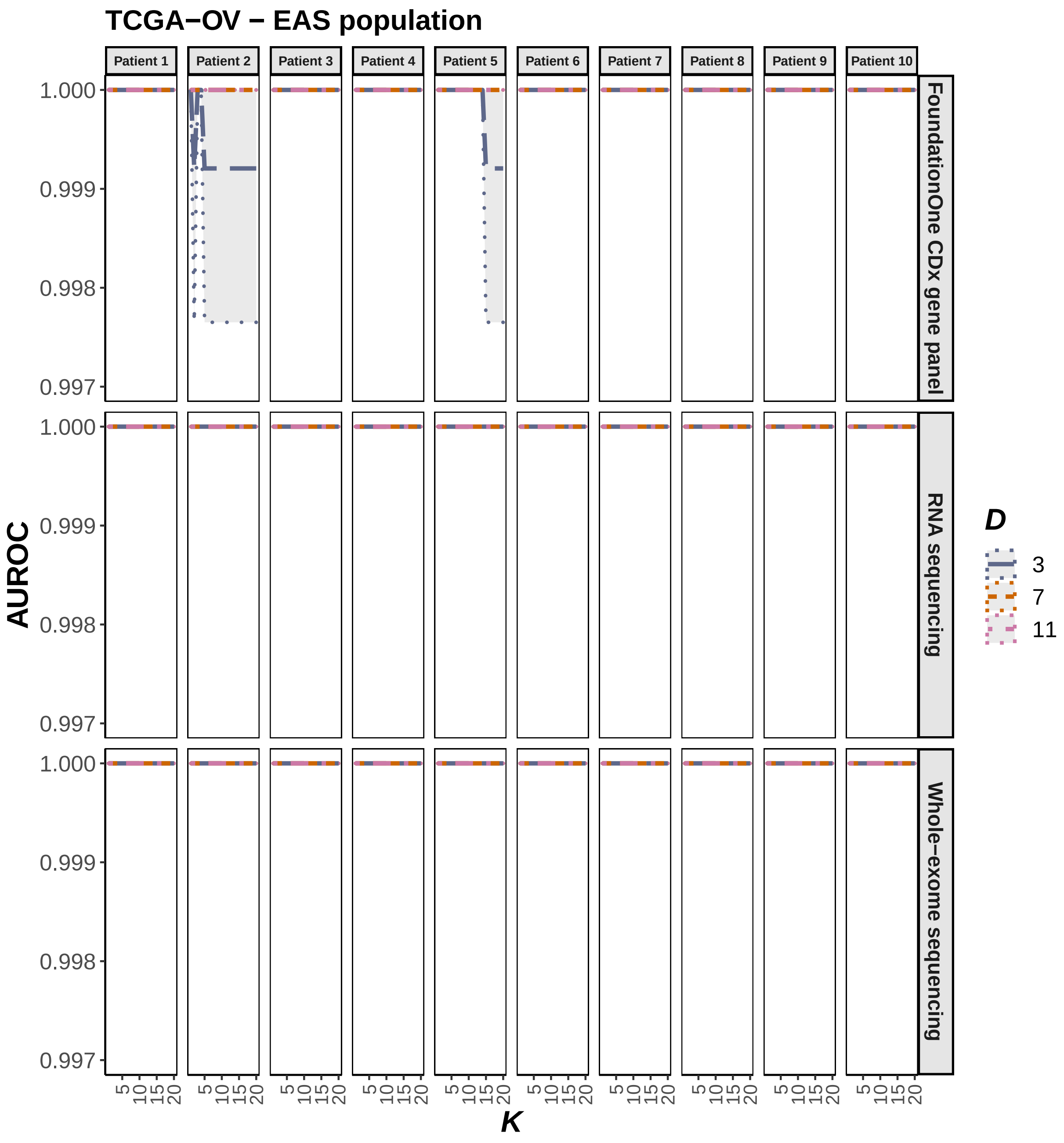

**Figure 1.** **E)** Dependence of EAS-specific AUROC on the inference parameters $D$ and $K$, computed using data synthesis for 10 TCGA-OV patients and the three profiling modalities: WES, RNA-seq and FoundationOne® CDx panels. The central AUROC values are shown in solid, and the 95% CI in dashed, lines.

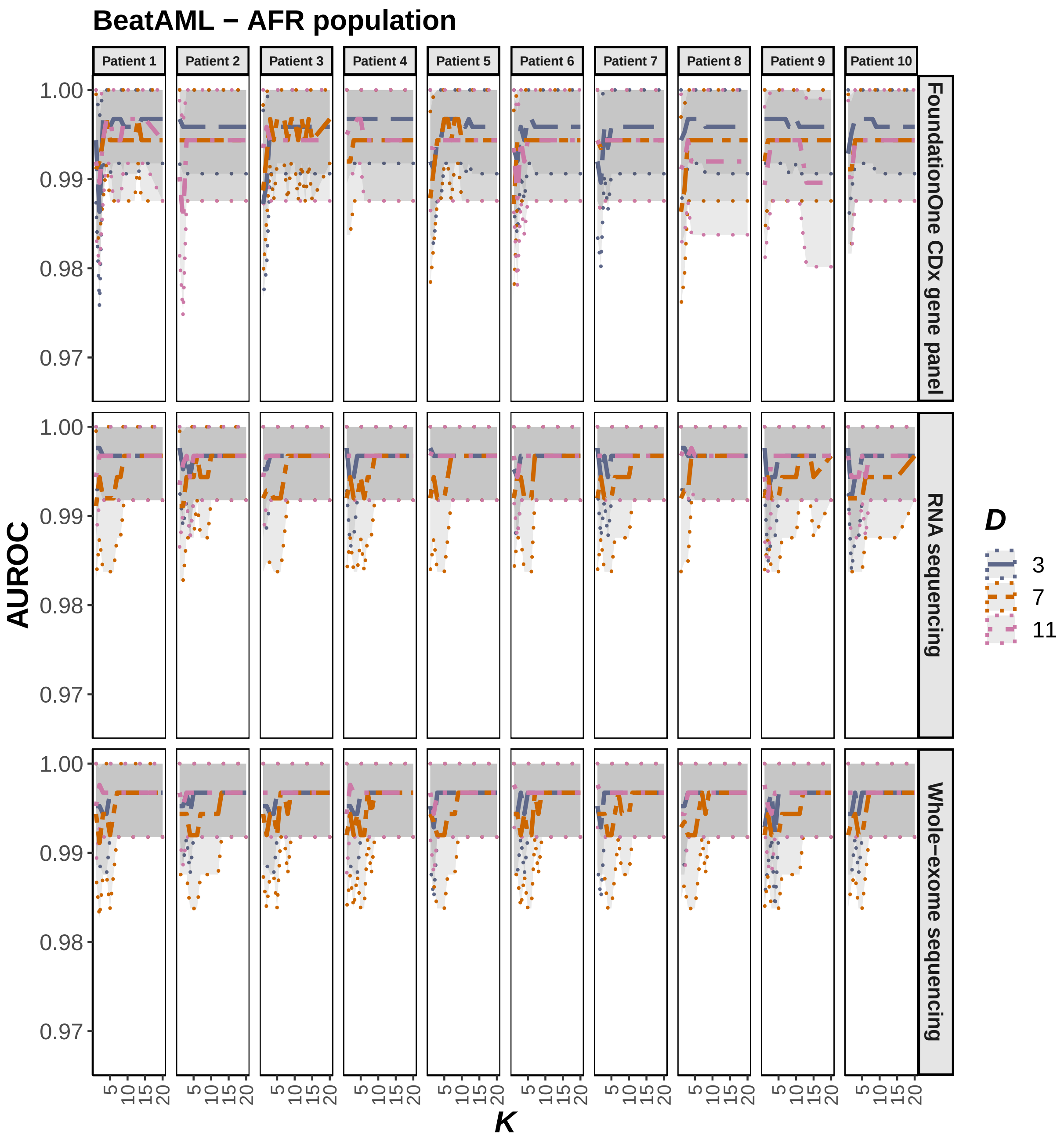

**Figure 1.** **F)** Dependence of AFR-specific AUROC on the inference parameters $D$ and $K$, computed using data synthesis for 10 Beat AML patients and the three profiling modalities: WES, RNA-seq and FoundationOne® CDx panels. The central AUROC values are shown in solid, and the 95% CI in dashed, lines.

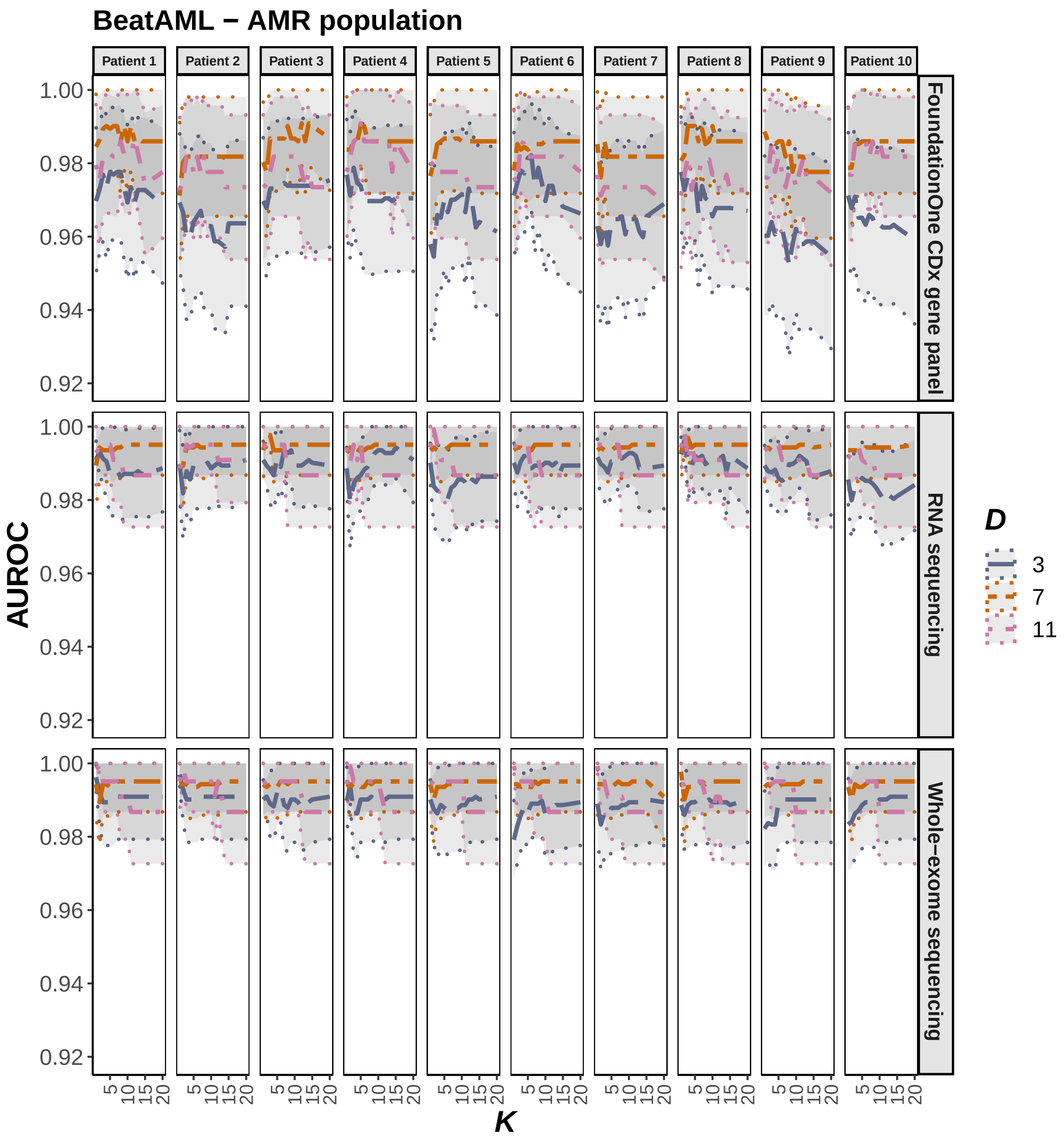

**Figure 1. G)** Dependence of AMR-specific AUROC on the inference parameters $D$ and $K$, computed using data synthesis for 10 Beat AML patients and the three profiling modalities: WES, RNA-seq and FoundationOne® CDx panels. The central AUROC values are shown in solid, and the 95% CI in dashed, lines.

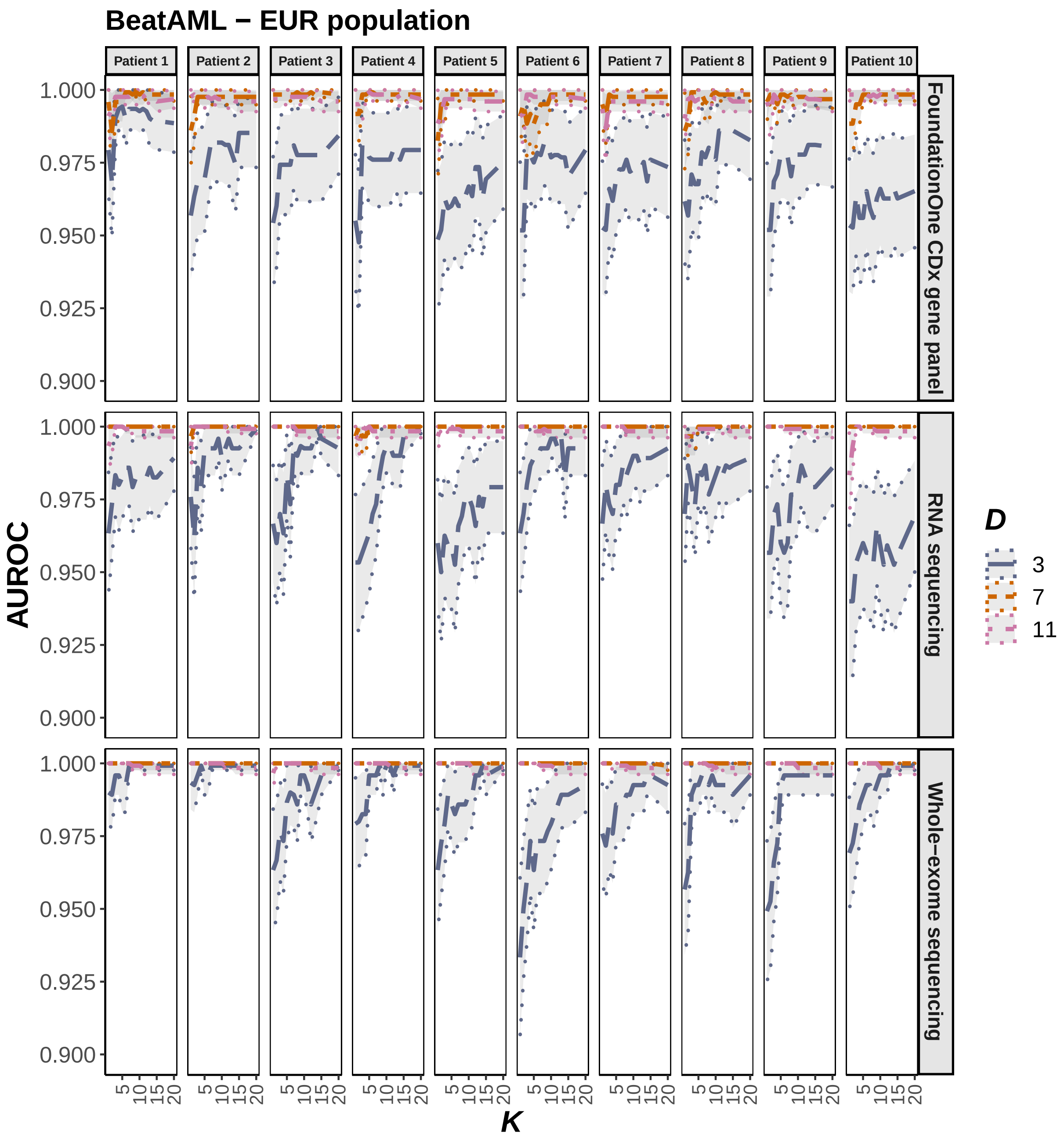

**Figure 1. H)** Dependence of EUR-specific AUROC on the inference parameters $D$ and $K$, computed using data synthesis for 10 Beat AML patients and the three profiling modalities: WES, RNA-seq and FoundationOne® CDx panels. The central AUROC values are shown in solid, and the 95% CI in dashed, lines.

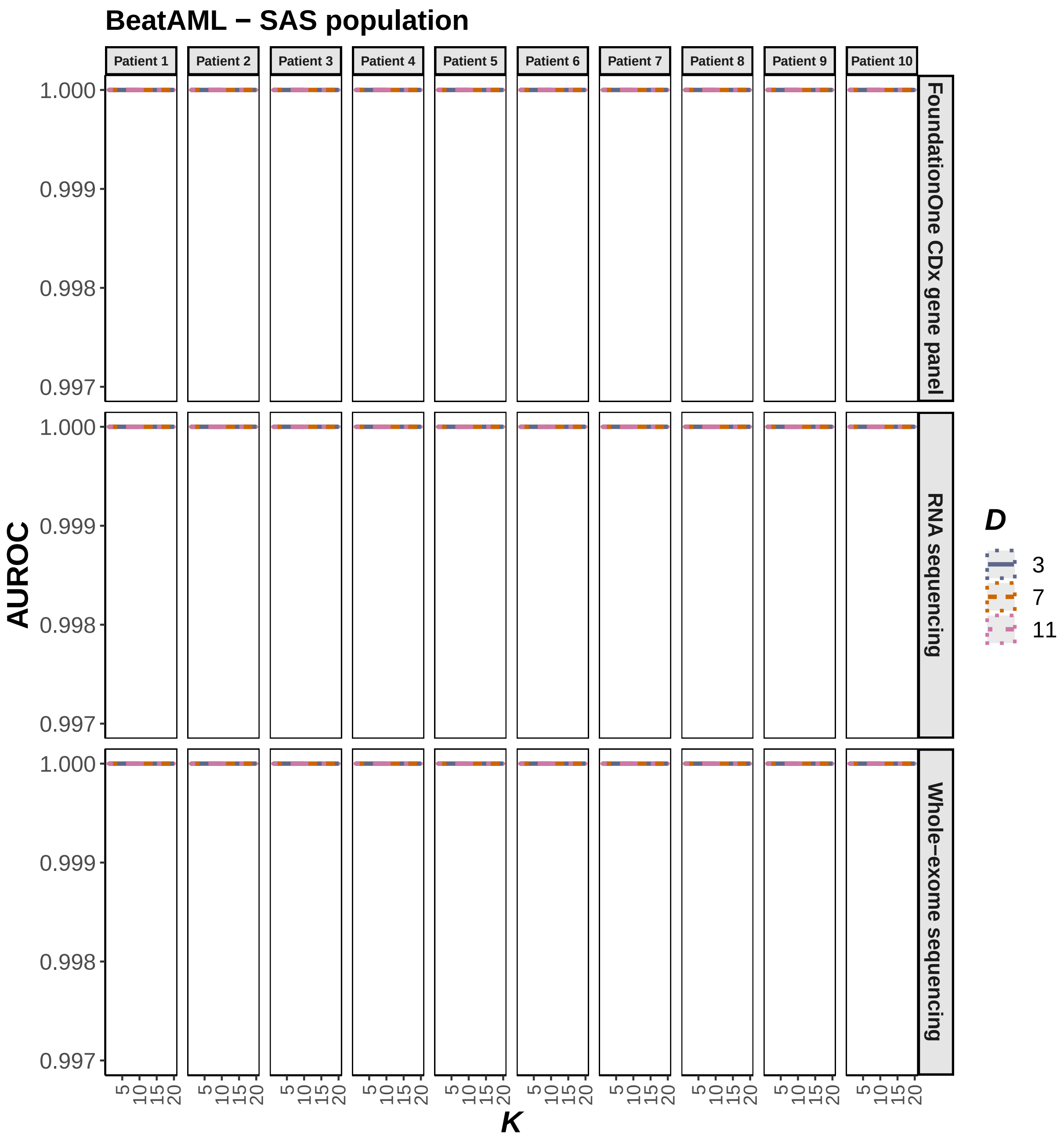

**Figure 1. I)** Dependence of SAS-specific AUROC on the inference parameters $D$ and $K$, computed using data synthesis for 10 Beat AML patients and the three profiling modalities: WES, RNA-seq and FoundationOne® CDx panels. The central AUROC values are shown in solid, and the 95% CI in dashed, lines.

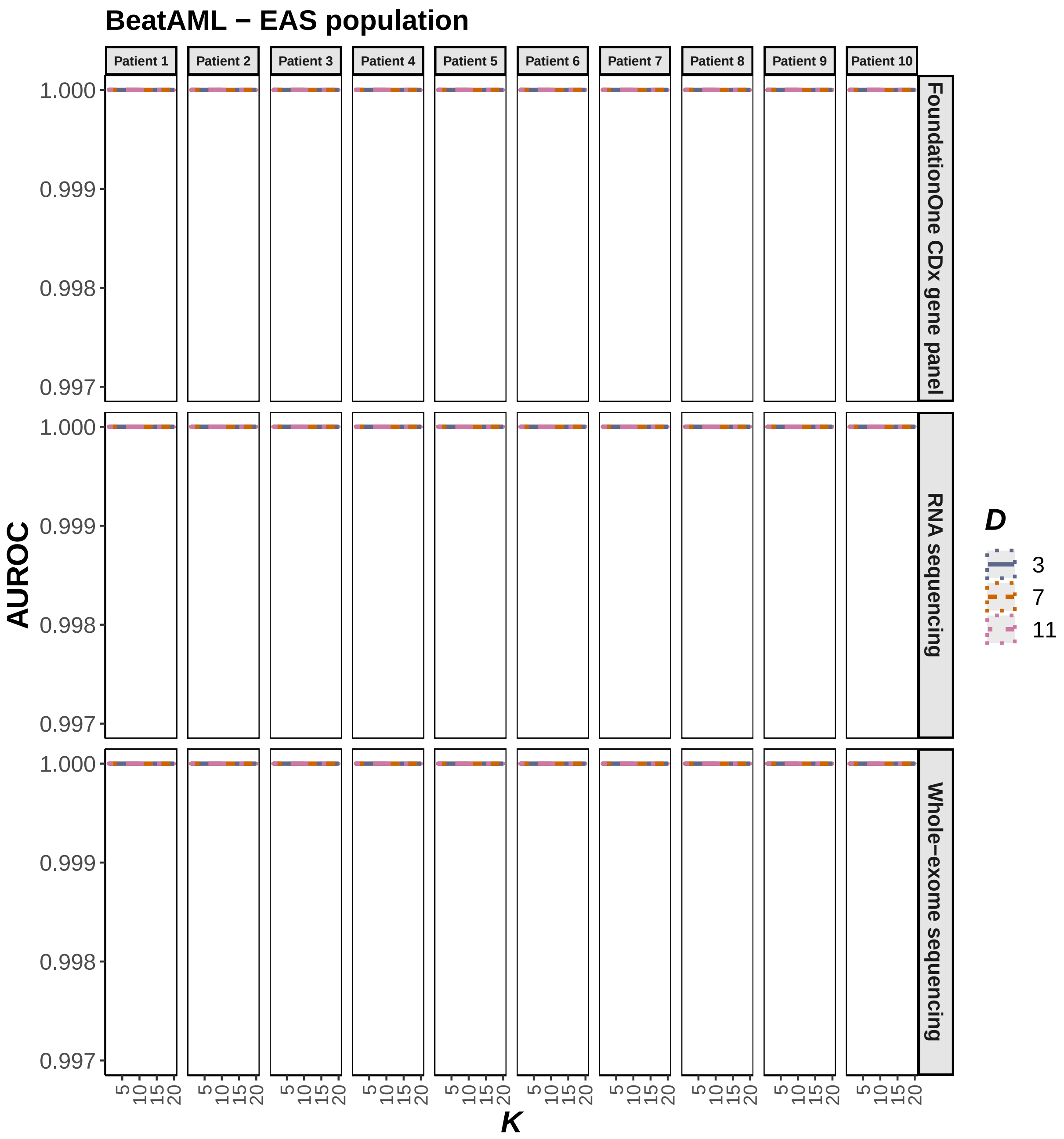

**Figure 1. J)** Dependence of EAS-specific AUROC on the inference parameters $D$ and $K$, computed using data synthesis for 10 Beat AML patients and the three profiling modalities: WES, RNA-seq and FoundationOne® CDx panels. The central AUROC values are shown in solid, and the 95% CI in dashed, lines.

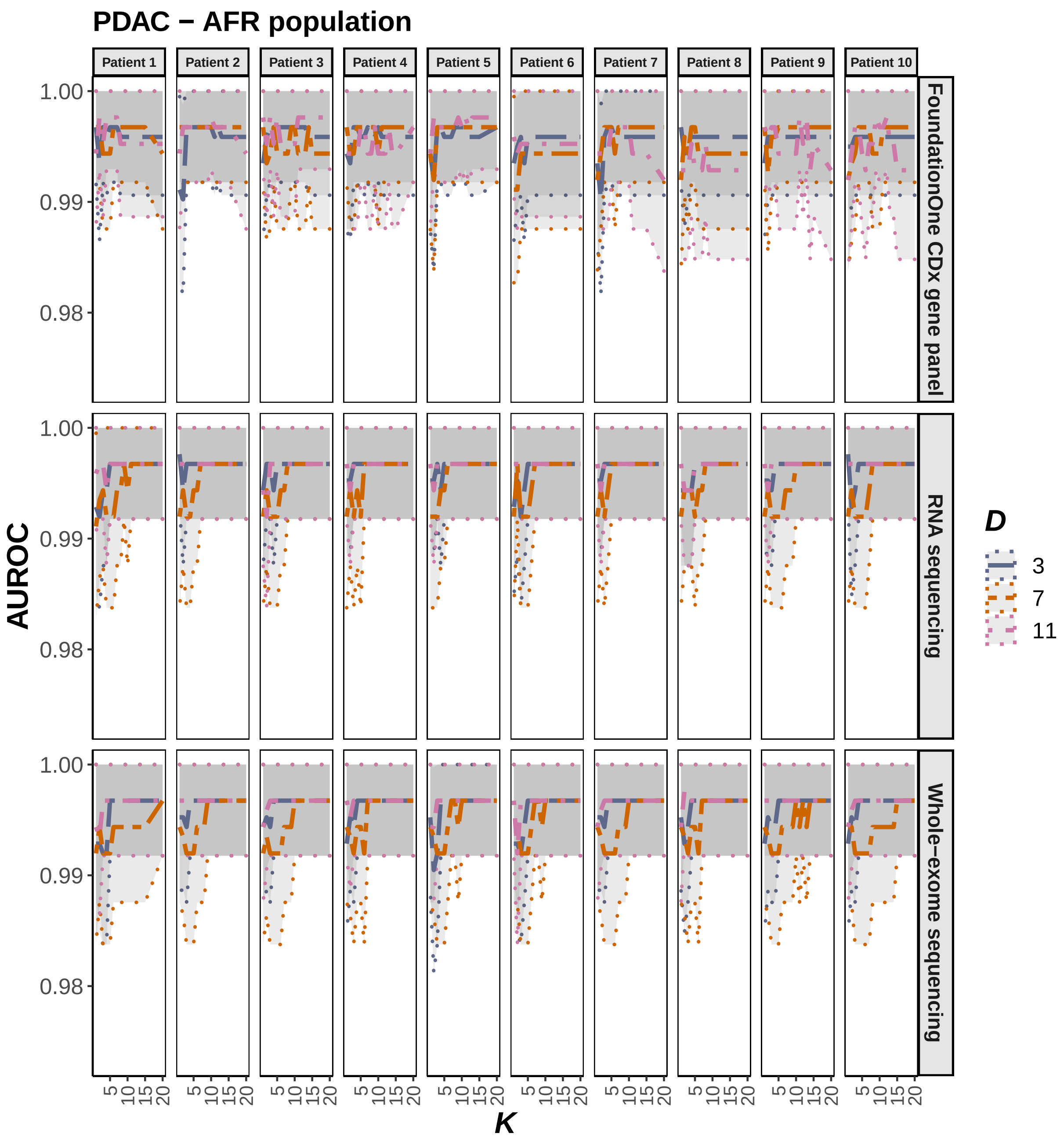

**Figure 1. K)** Dependence of AFR-specific AUROC on the inference parameters $D$ and $K$, computed using data synthesis for 10 PDAC patients and the three profiling modalities: WES, RNA-seq and FoundationOne® CDx panels. The central AUROC values are shown in solid, and the 95% CI in dashed, lines.

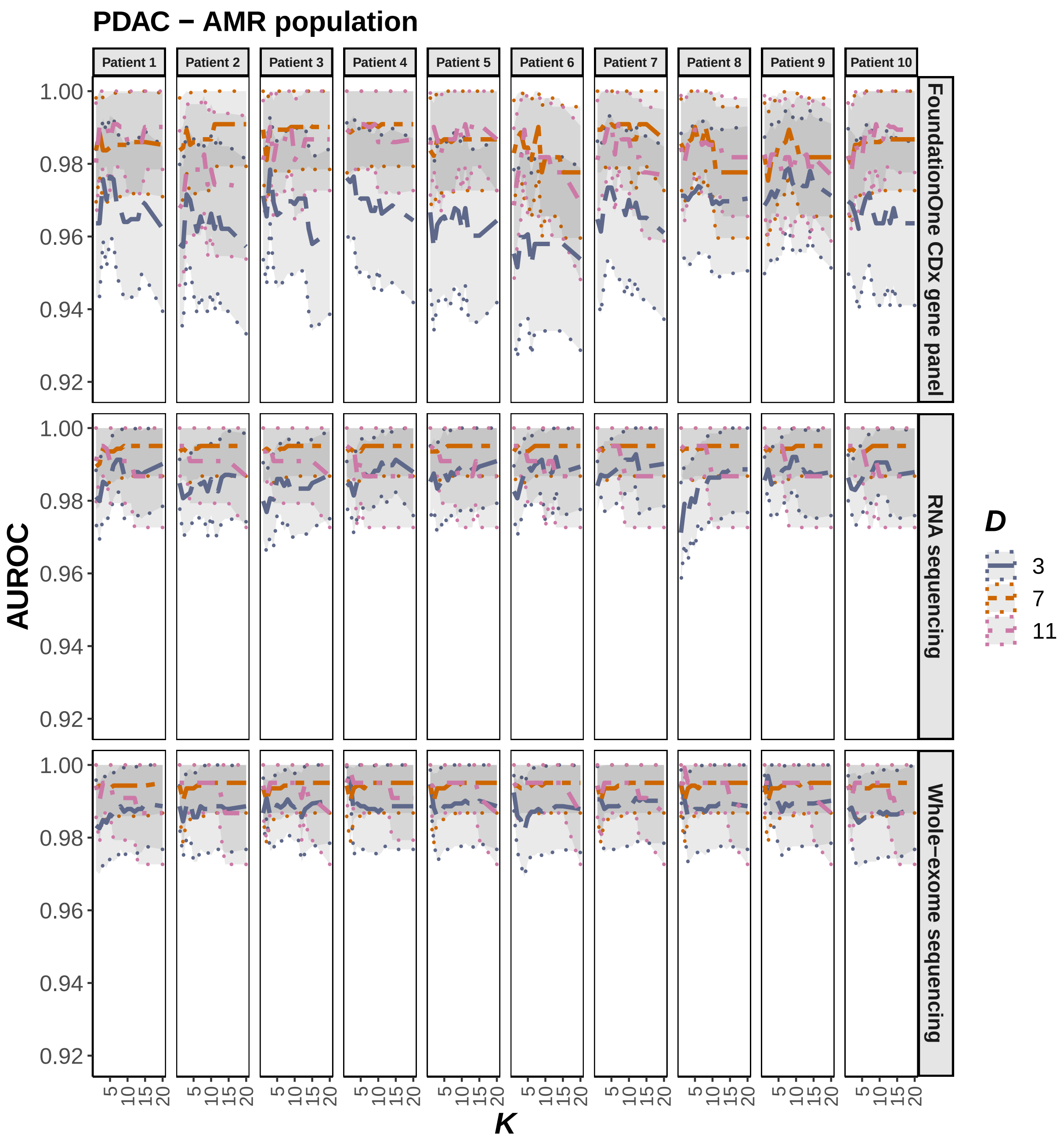

**Figure 1. L)** Dependence of AMR-specific AUROC on the inference parameters $D$ and $K$, computed using data synthesis for 10 PDAC patients and the three profiling modalities: WES, RNA-seq and FoundationOne® CDx panels. The central AUROC values are shown in solid, and the 95% CI in dashed, lines.

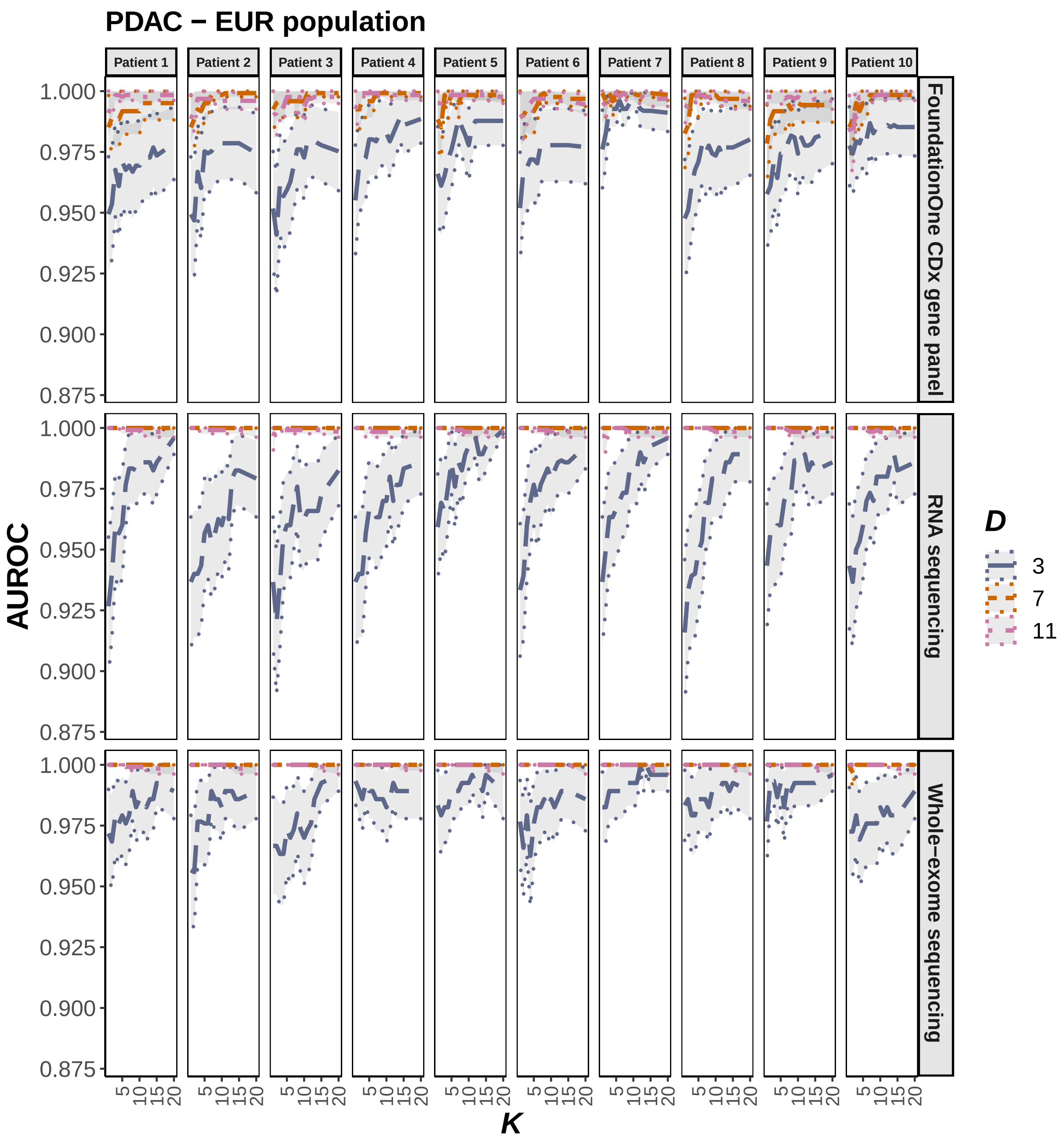

**Figure 1. M)** Dependence of EUR-specific AUROC on the inference parameters $D$ and $K$, computed using data synthesis for 10 PDAC patients and the three profiling modalities: WES, RNA-seq and FoundationOne® CDx panels. The central AUROC values are shown in solid, and the 95% CI in dashed, lines.

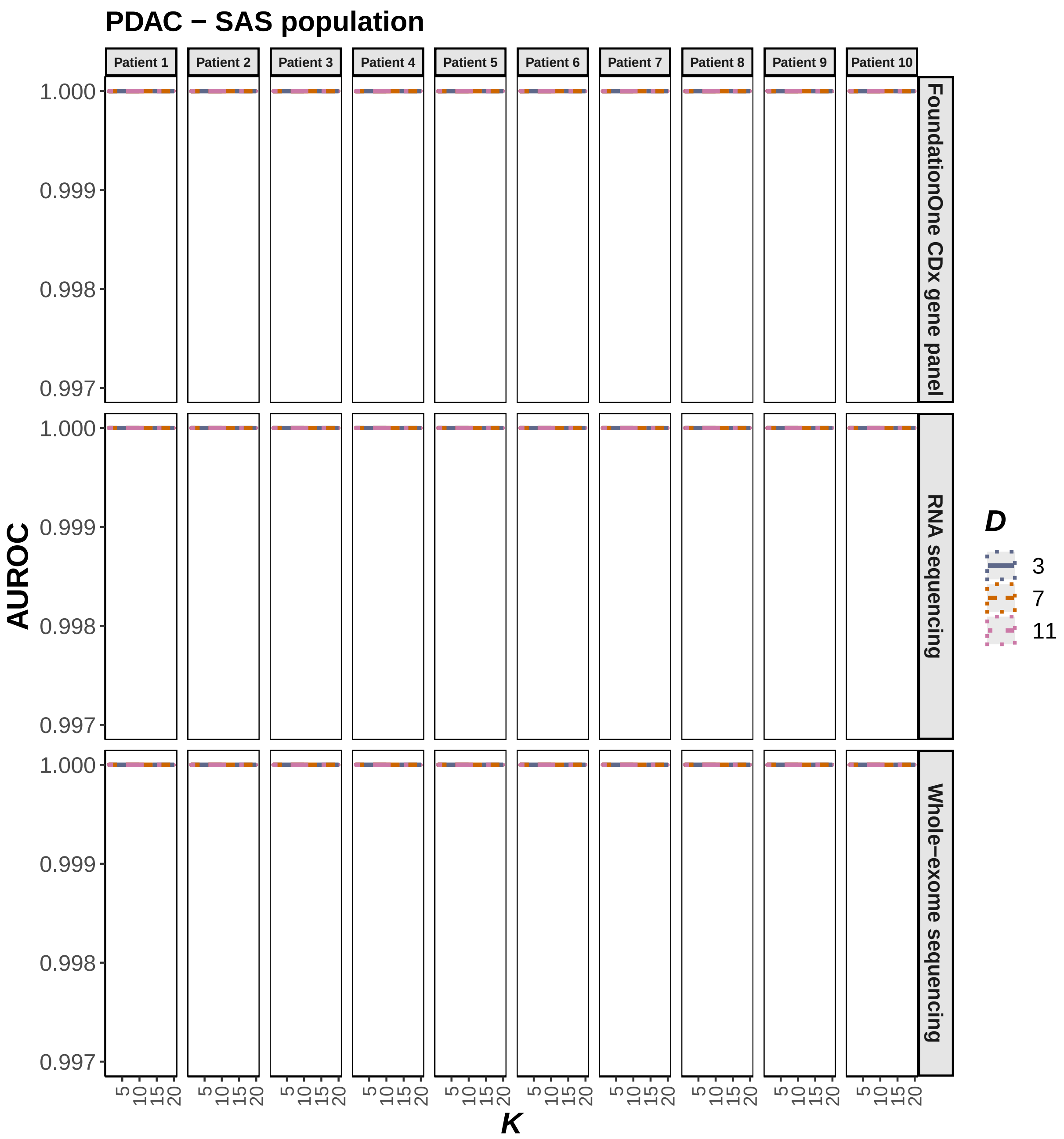

**Figure 1. N)** Dependence of SAS-specific AUROC on the inference parameters $D$ and $K$, computed using data synthesis for 10 PDAC patients and the three profiling modalities: WES, RNA-seq and FoundationOne® CDx panels. The central AUROC values are shown in solid, and the 95% CI in dashed, lines.

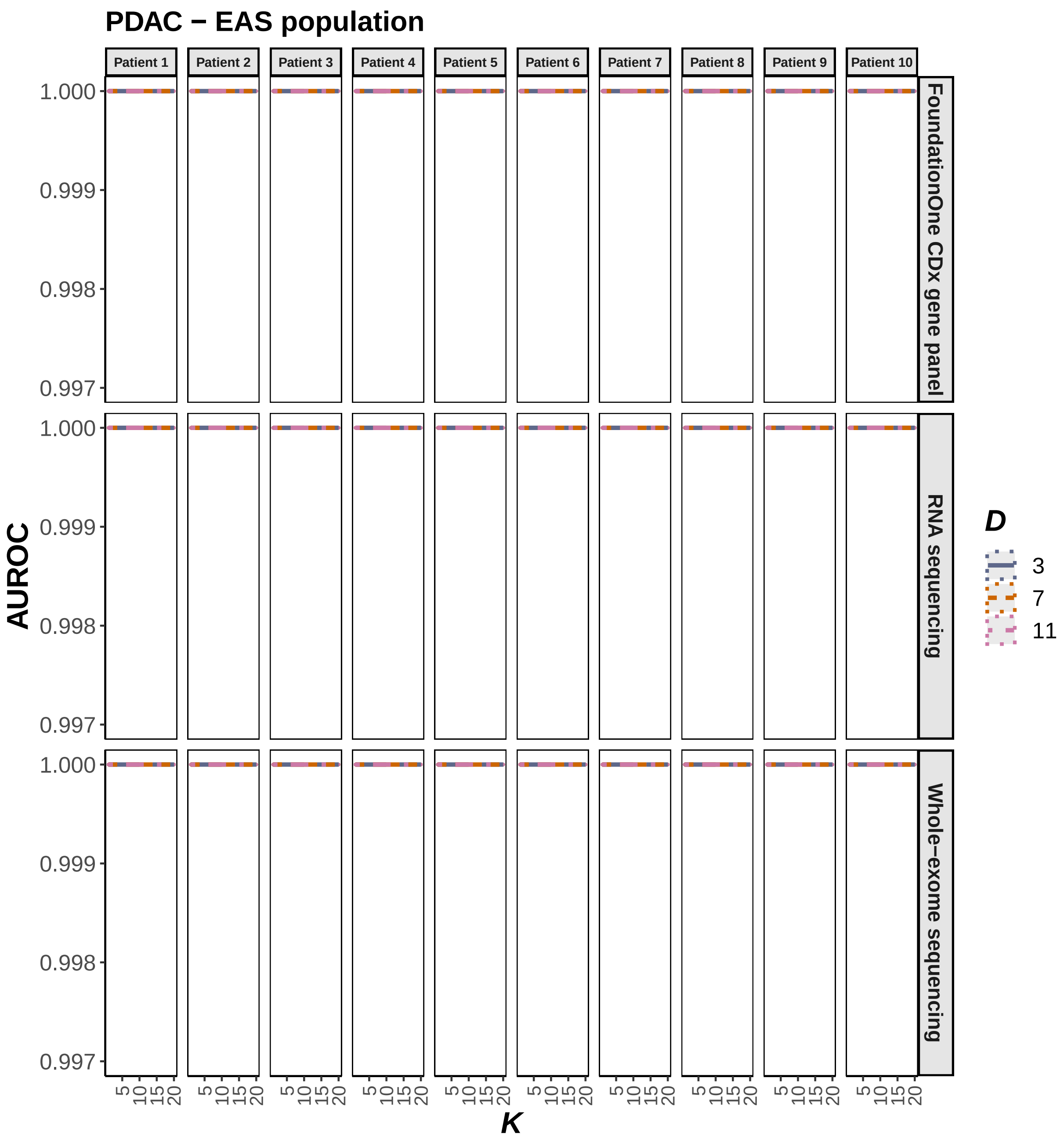

**Figure 1. O)** Dependence of EAS-specific AUROC on the inference parameters $D$ and $K$, computed using data synthesis for 10 PDAC patients and the three profiling modalities: WES, RNA-seq and FoundationOne® CDx panels. The central AUROC values are shown in solid, and the 95% CI in dashed, lines.

### Performance of genetic ancestry inference as a function of coverage

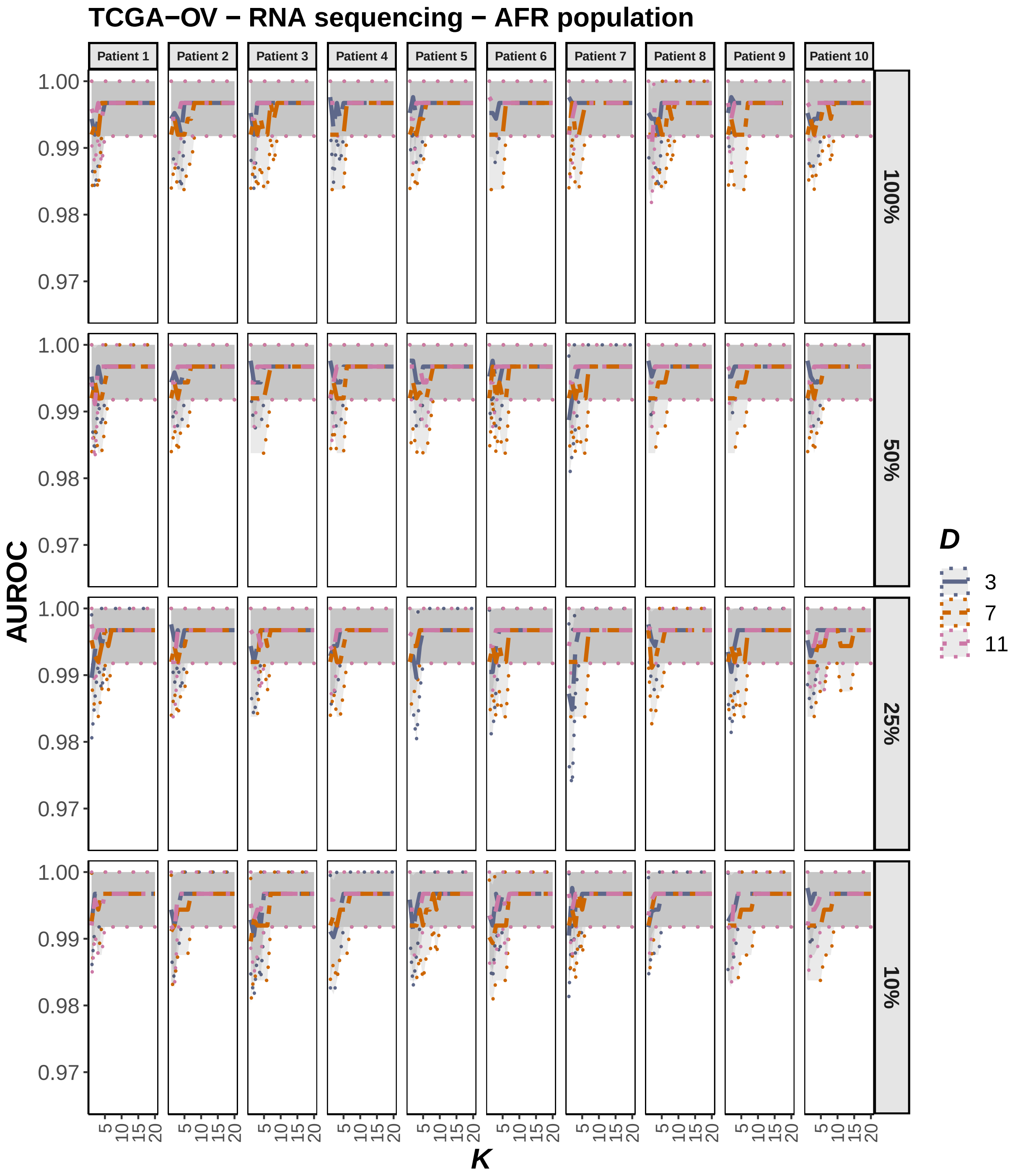

**Figure 2. A)** Dependence of AFR-specific AUROC on the inference parameters $D$ and $K$, at the original and reduced sequence coverage values, as indicated by the percentages of the original coverage. AUROC was computed using data synthesis for RNA-seq profiles of 10 TCGA-OV patients. The central AUROC values are shown in solid, and the 95% CI in dashed, lines.

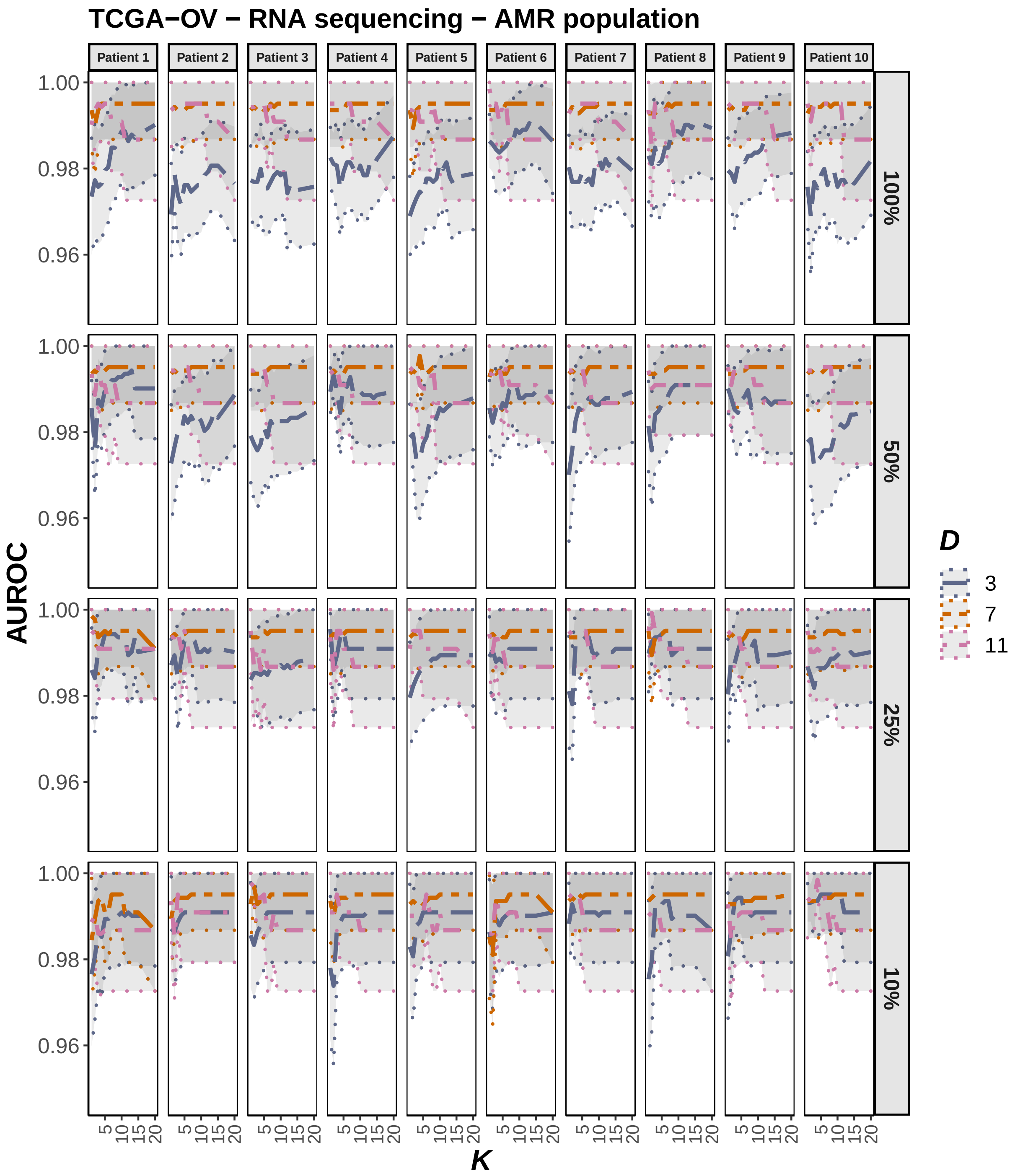

**Figure 2. B)** Dependence of AMR-specific AUROC on the inference parameters $D$ and $K$, at the original and reduced sequence coverage values, as indicated by the percentages of the original coverage. AUROC was computed using data synthesis for RNA-seq profiles of 10 TCGA-OV patients. The central AUROC values are shown in solid, and the 95% CI in dashed, lines.

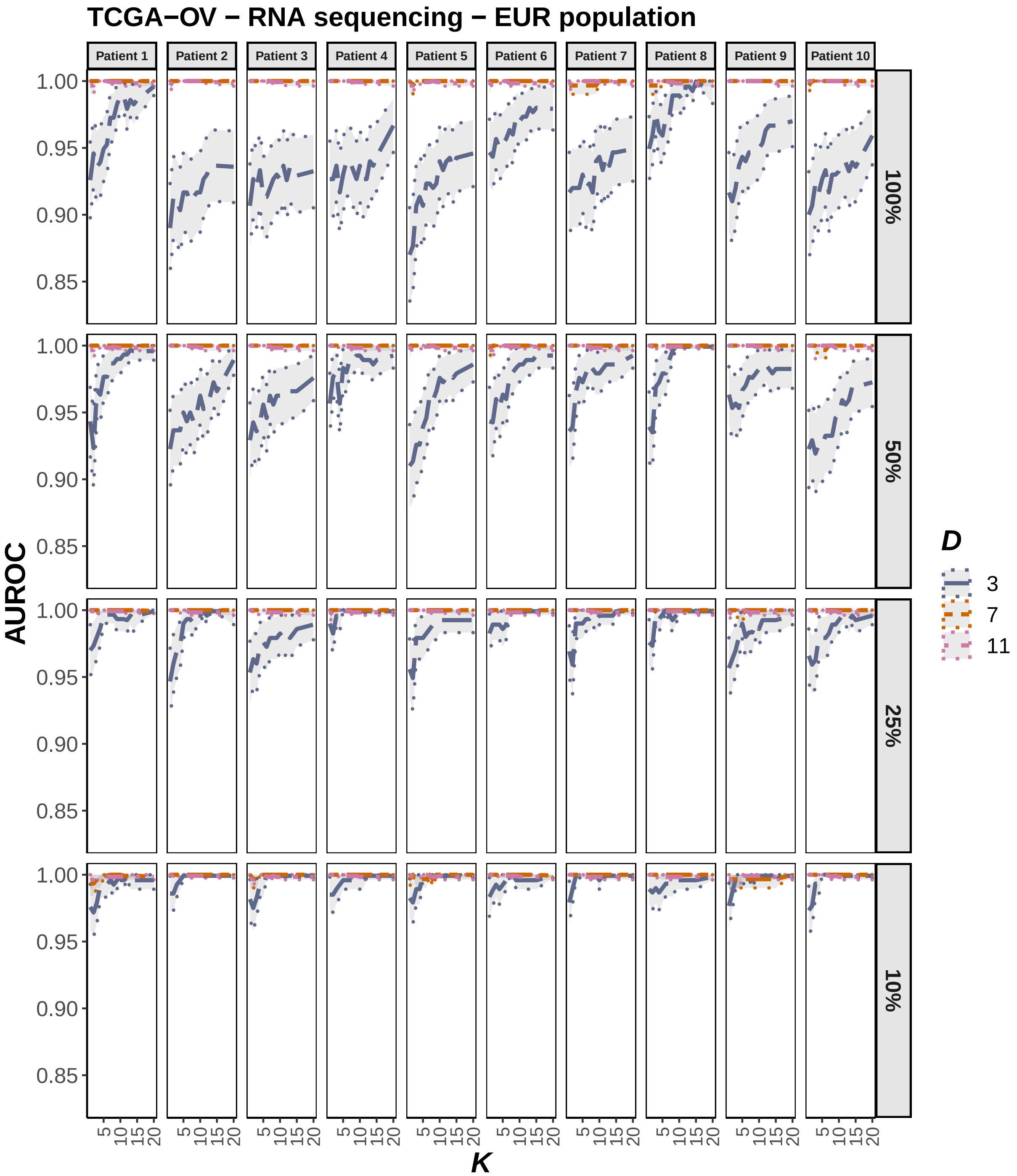

**Figure 2. C)** Dependence of EUR-specific AUROC on the inference parameters $D$ and $K$, at the original and reduced sequence coverage values, as indicated by the percentages of the original coverage. AUROC was computed using data synthesis for RNA-seq profiles of 10 TCGA-OV patients. The central AUROC values are shown in solid, and the 95% CI in dashed, lines.

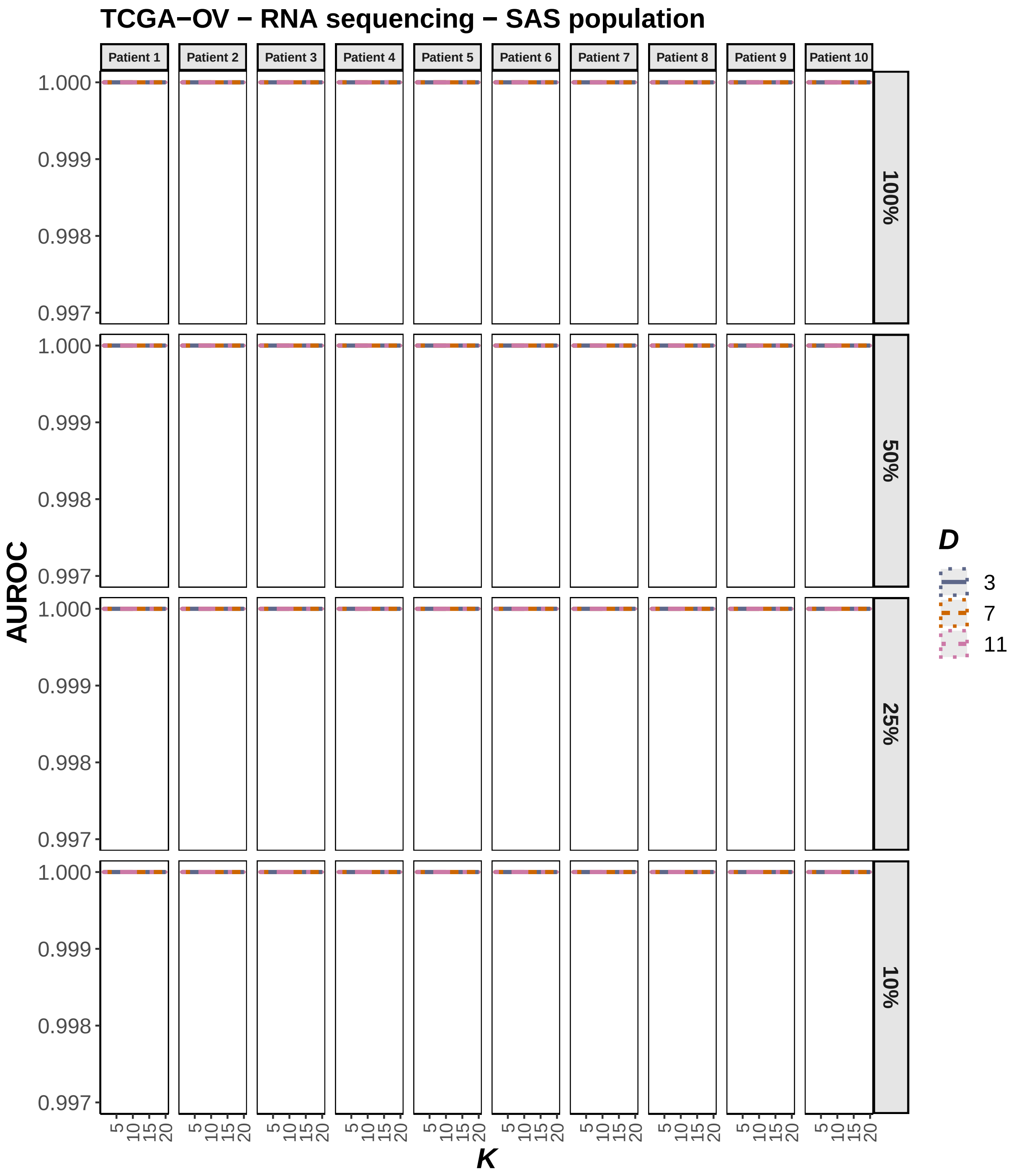

**Figure 2. D)** Dependence of SAS-specific AUROC on the inference parameters $D$ and $K$, at the original and reduced sequence coverage values, as indicated by the percentages of the original coverage. AUROC was computed using data synthesis for RNA-seq profiles of 10 TCGA-OV patients. The central AUROC values are shown in solid, and the 95% CI in dashed, lines.

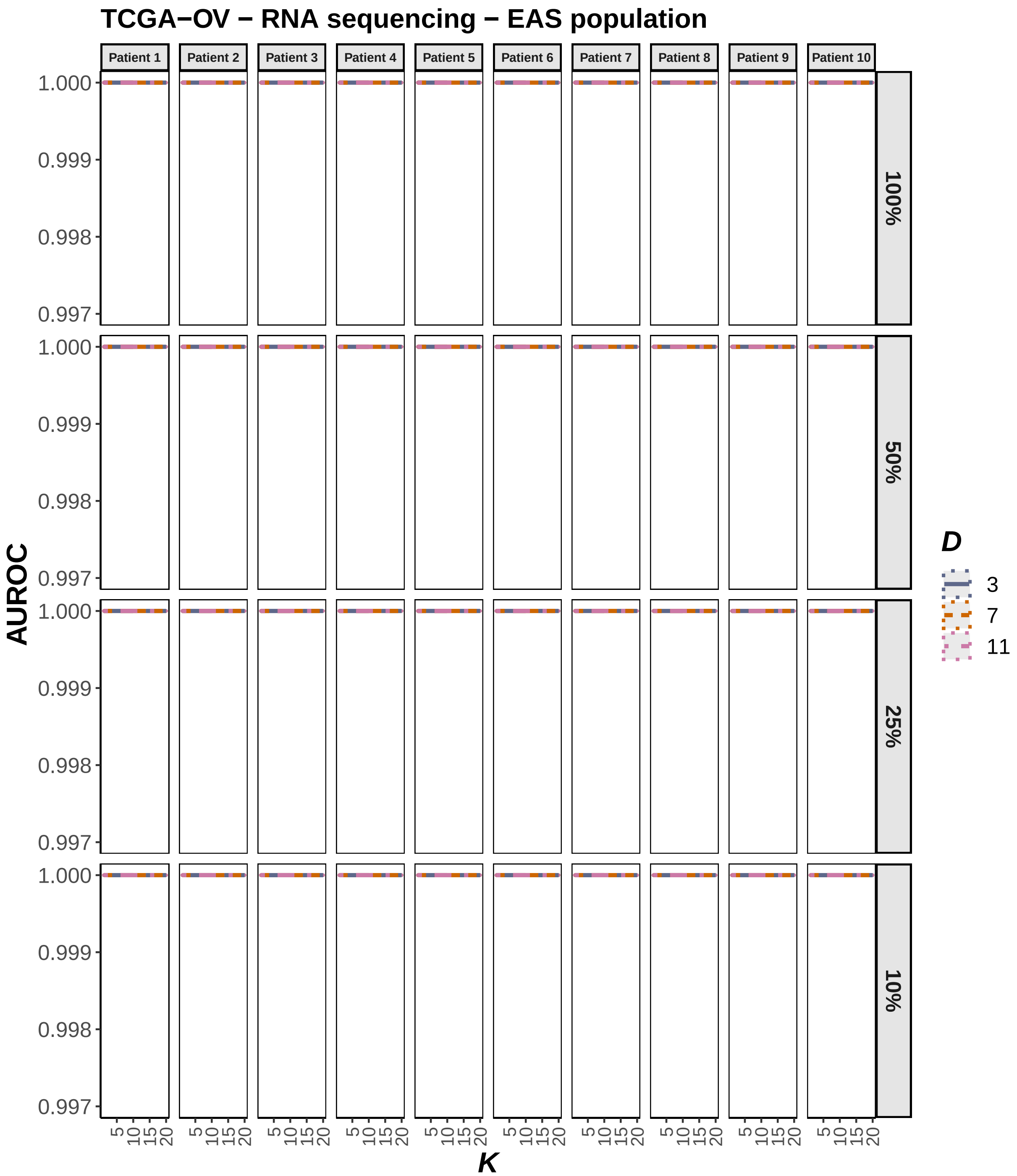

**Figure 2. E)** Dependence of EAS-specific AUROC on the inference parameters $D$ and $K$, at the original and reduced sequence coverage values, as indicated by the percentages of the original coverage. AUROC was computed using data synthesis for RNA-seq profiles of 10 TCGA-OV patients. The central AUROC values are shown in solid, and the 95% CI in dashed, lines.

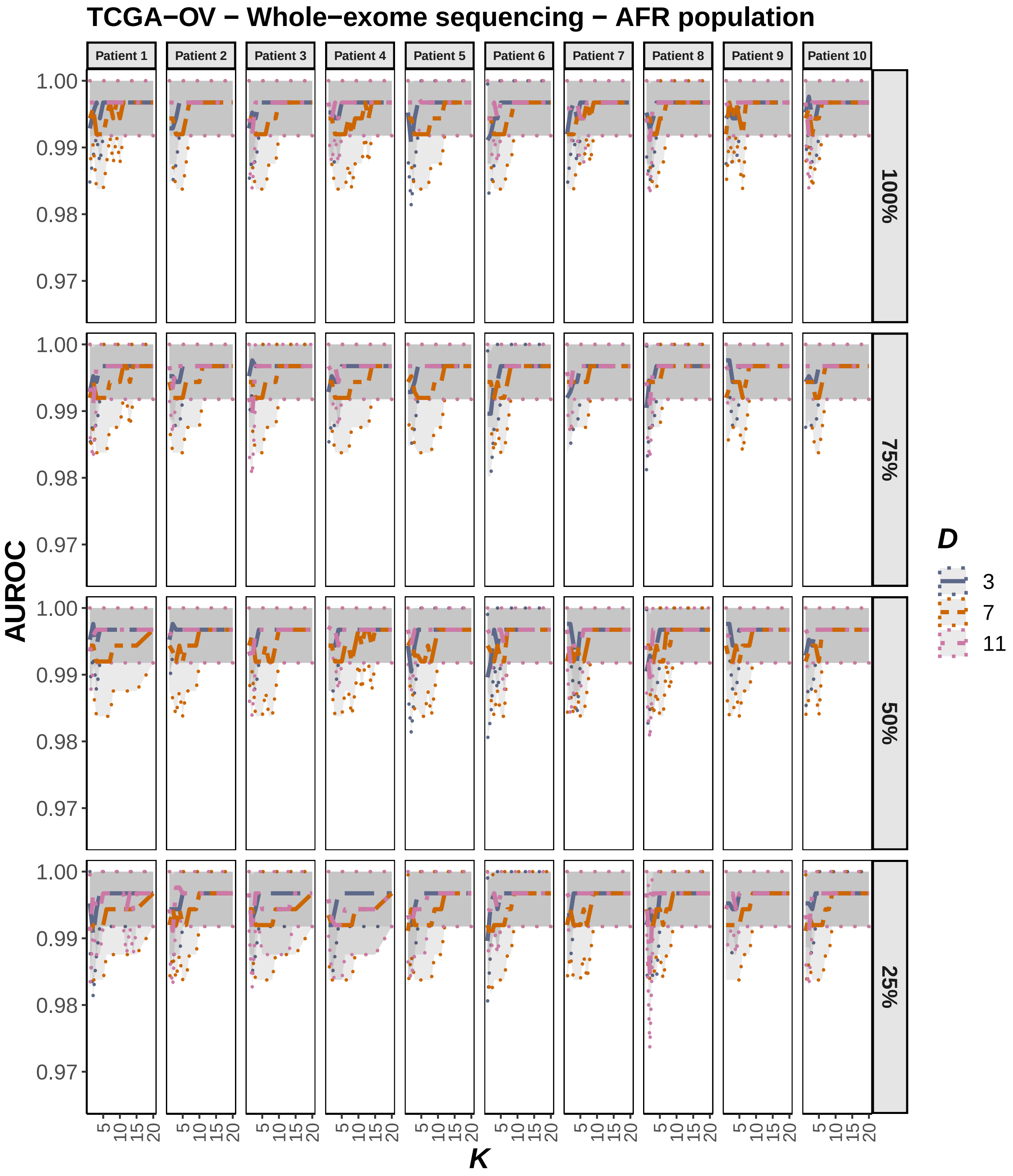

**Figure 2. F)** Dependence of AFR-specific AUROC on the inference parameters $D$ and $K$, at the original and reduced sequence coverage values, as indicated by the percentages of the original coverage. AUROC was computed using data synthesis for WES profiles of 10 TCGA-OV patients. The central AUROC values are shown in solid, and the 95% CI in dashed, lines.

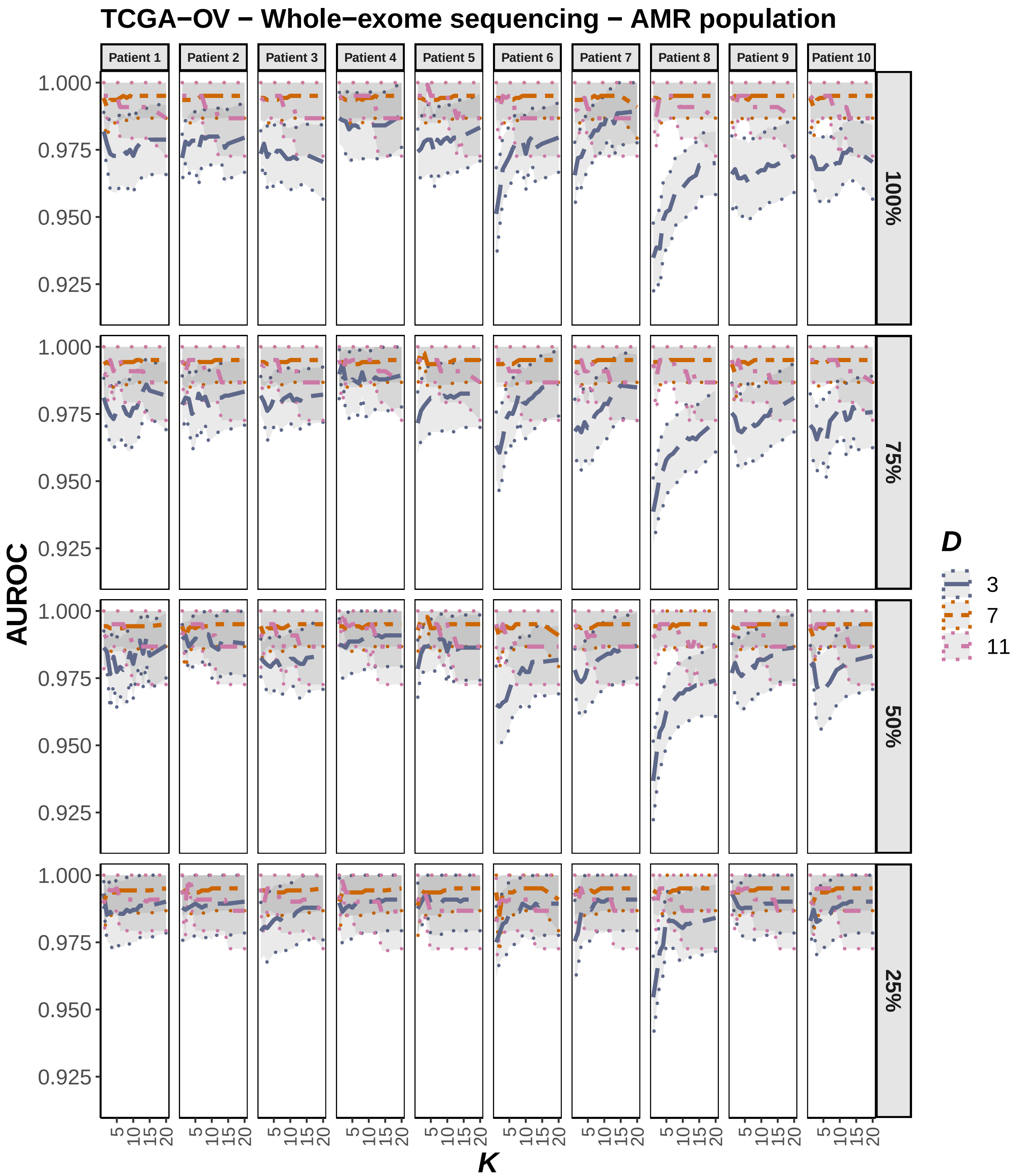

**Figure 2. G)** Dependence of AMR-specific AUROC on the inference parameters $D$ and $K$, at the original and reduced sequence coverage values, as indicated by the percentages of the original coverage. AUROC was computed using data synthesis for WES profiles of 10 TCGA-OV patients. The central AUROC values are shown in solid, and the 95% CI in dashed, lines.

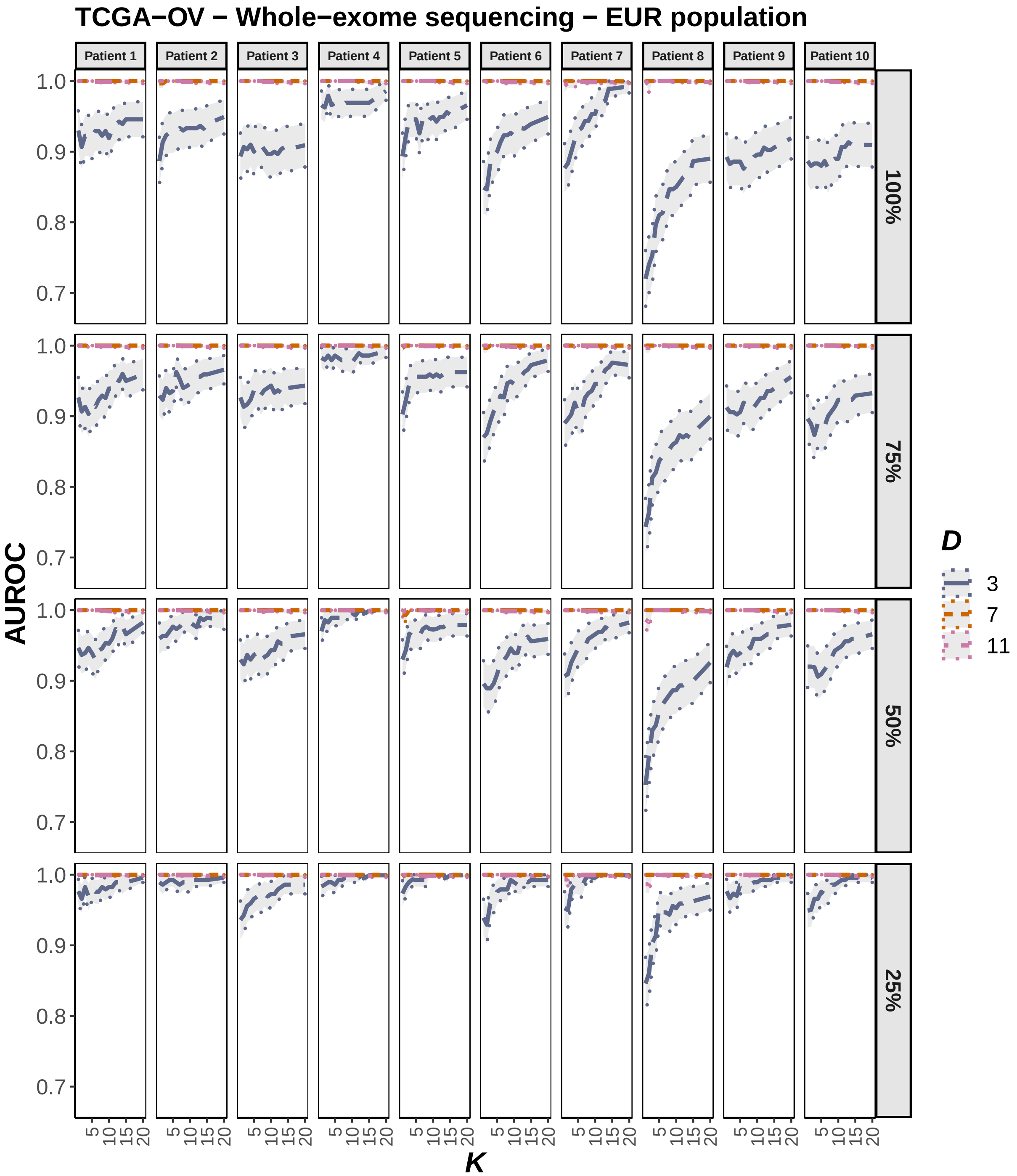

**Figure 2.** **H)** Dependence of EUR-specific AUROC on the inference parameters $D$ and $K$, at the original and reduced sequence coverage values, as indicated by the percentages of the original coverage. AUROC was computed using data synthesis for WES profiles of 10 TCGA-OV patients. The central AUROC values are shown in solid, and the 95% CI in dashed, lines.

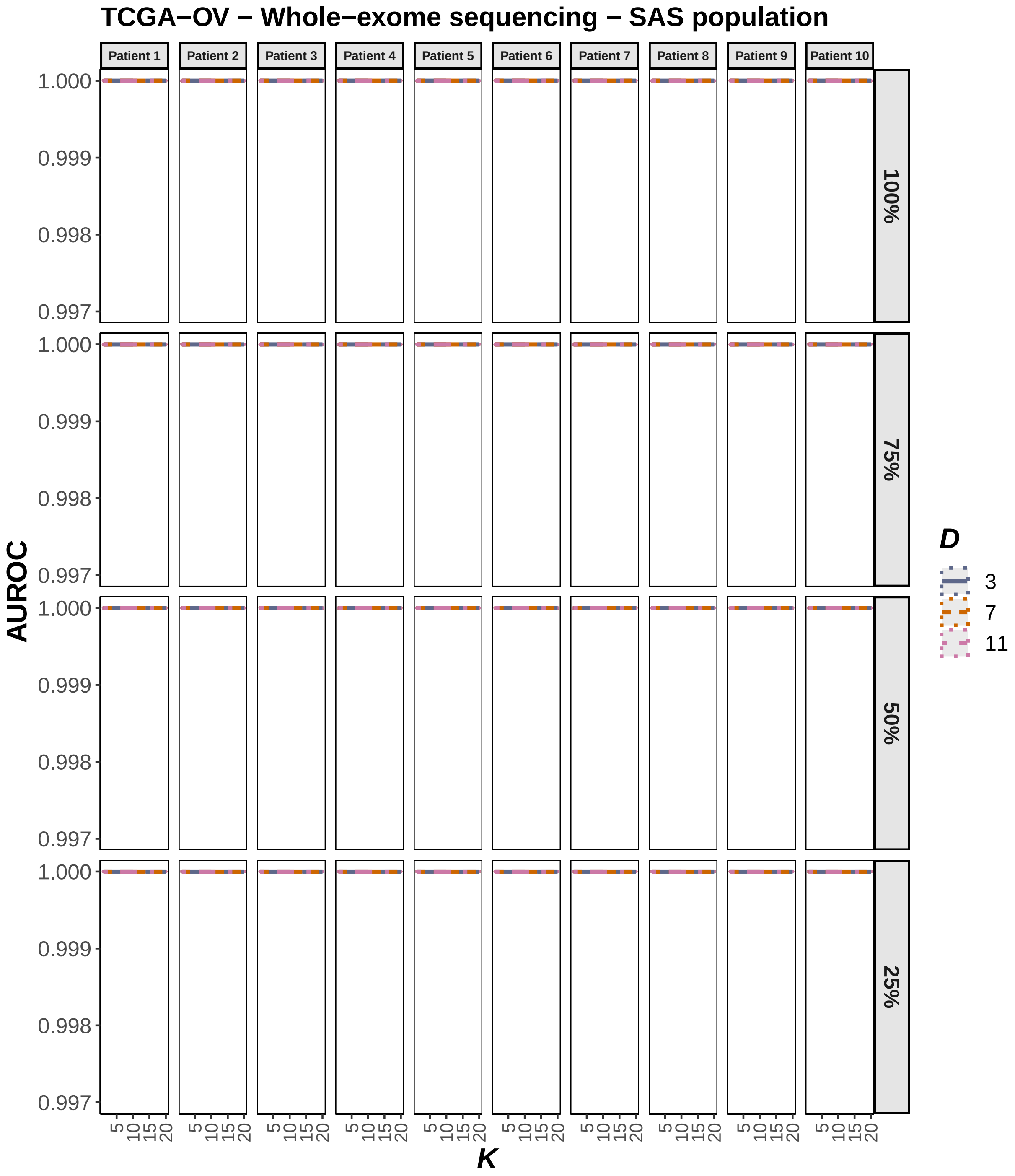

**Figure 2. I)** Dependence of SAS-specific AUROC on the inference parameters $D$ and $K$, at the original and reduced sequence coverage values, as indicated by the percentages of the original coverage. AUROC was computed using data synthesis for WES profiles of 10 TCGA-OV patients. The central AUROC values are shown in solid, and the 95% CI in dashed, lines.

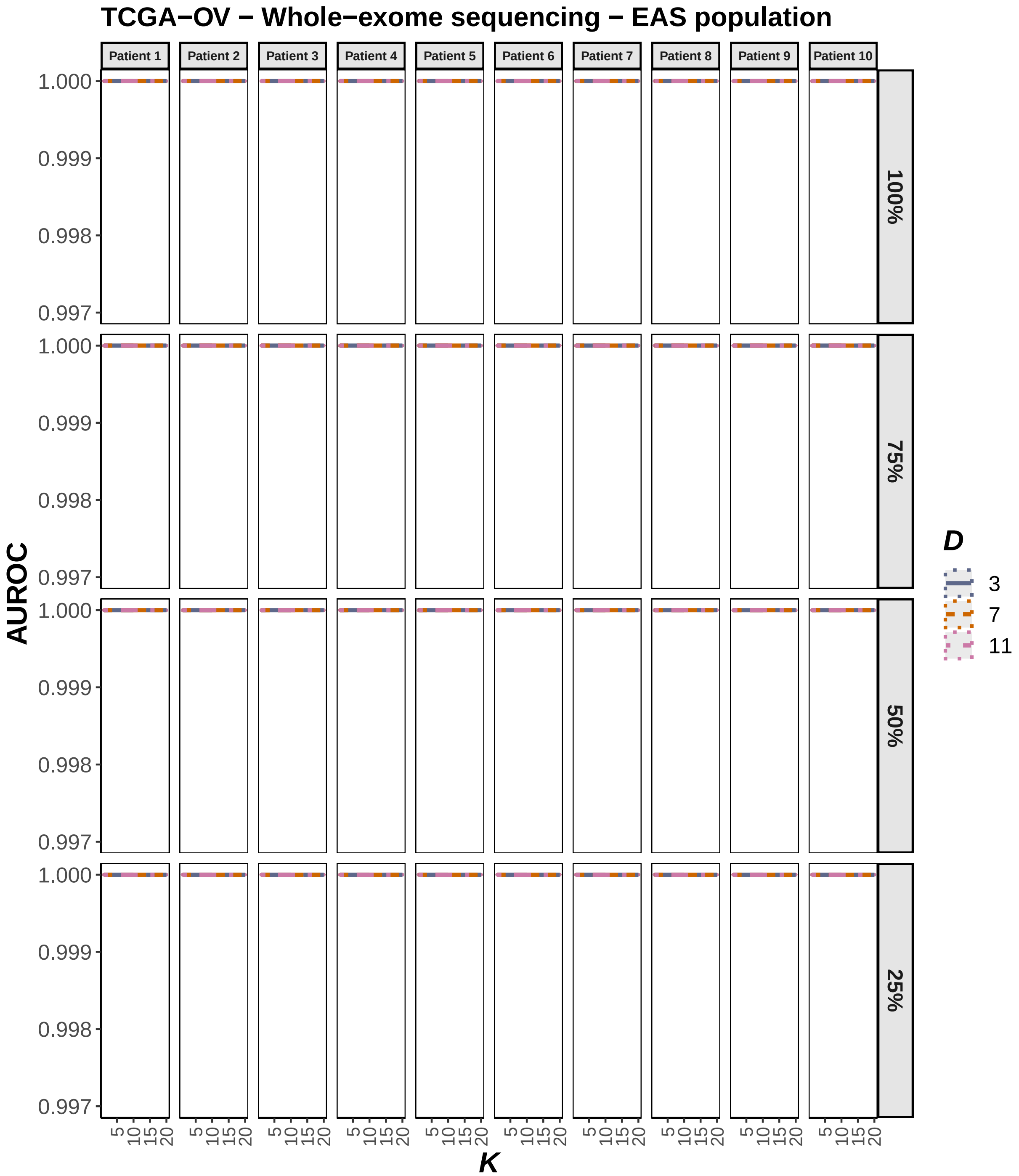

**Figure 2. J)** Dependence of EAS-specific AUROC on the inference parameters $D$ and $K$, at the original and reduced sequence coverage values, as indicated by the percentages of the original coverage. AUROC was computed using data synthesis for WES profiles of 10 TCGA-OV patients. The central AUROC values are shown in solid, and the 95% CI in dashed, lines.

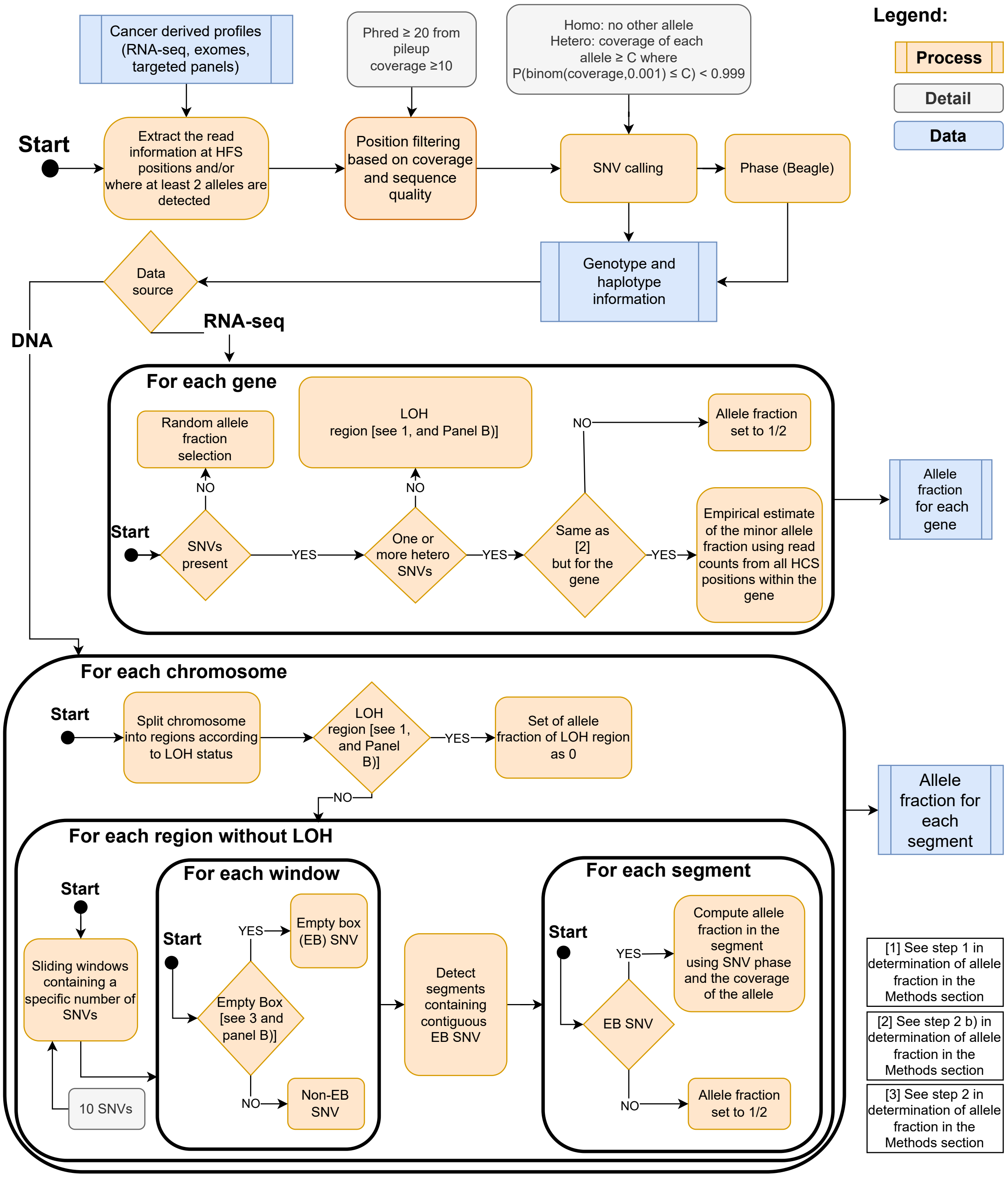

**Figure 3. A)** Overview of the procedure for allele fraction estimation in genes and segments.

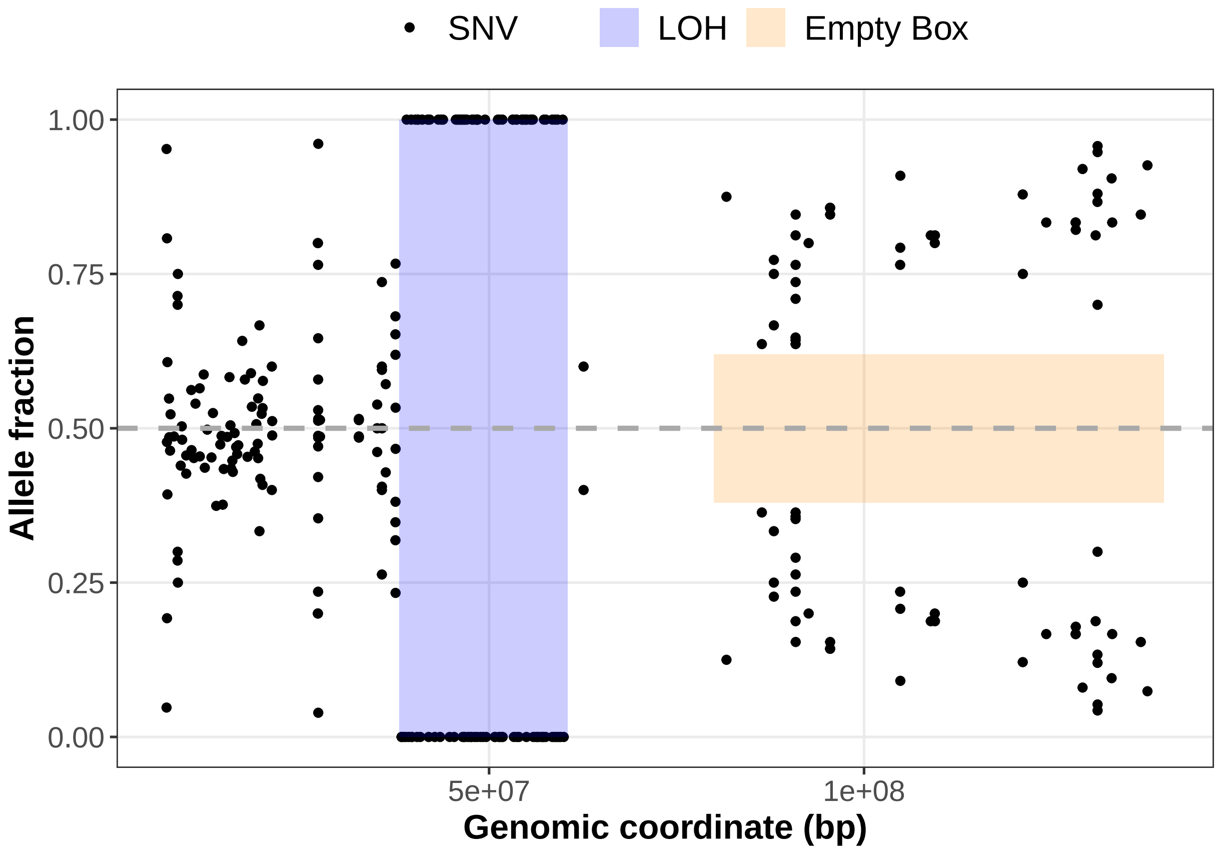

**Figure 3. B)** Estimation of the allele fractions illustrating the key concepts used. Each data point represents an observed variant allele fraction at the corresponding genomic coordinate.
